# Single-cell mapping of the fallopian tube reveals a genomically unstable secretory cell state enriched in carriers of germline *BRCA1/2* mutations

**DOI:** 10.64898/2026.08.26.746998

**Authors:** Yaniv Eyal-Lubling, Maria Vias, Katarzyna Kania, Dimana Kaludova, James Hall, Robin Crawford, Rutendo Nyagumbo, Sophie Ward, Svetlana Khoronenkova, Samuel Aparicio, Charles Swanton, Mercedes Jimenez Linan, James D. Brenton, Filipe Correia Martins

**Author notes:** Contributed equally (Yaniv Eyal-Lubling, Maria Vias). **This pdf has been formatted for convenient screen reading.** Citations and figures have active links indicated by coloured text. After following a link, pdf browsers will navigate back to the original page with the “go back to previous view” button or keyboard short-cuts (e.g. Alt/Command + left arrow).

## Abstract

Carriers of germline *BRCA1* or *BRCA2* alterations have a substantially increased lifetime risk of high-grade serous ovarian carcinoma (HGSOC), which originates from the secretory cells of the fallopian tube. However, comparative multi-omic analyses of bulk fallopian tube tissue from *BRCA1/2* carriers and the general population have, to date, revealed only limited differences. New molecular biomarkers of early malignant transformation in the FT are needed to enable non-surgical cancer interception in high-risk individuals through window-of-opportunity trials prior to risk-reducing surgery.

We performed a comprehensive single-cell, multi-regional analysis of fallopian tubes from 34 women, including 15 carriers of germline *BRCA1/2* alterations. Using a metacell-based approach applied to single-cell transcriptomic data, we identify both established and previously unrecognised cellular populations, and characterise phenotypic variation associated with menopausal status, menstrual cycle phase, hormonal contraception use, and anatomical region of the fallopian tube.

Menopause was associated with depletion of ciliated cells, whilst both secretory (SEC) and ciliated epithelial cells (CEC) shifted to a glandular phenotype in the luteal phase. Previous hormonal contraception usage had lasting effects including depletion of CD163-positive tissue resident macrophages and progesterone-specific increase of MHC-II expression in SECs.

Metacell analysis further identified distinct subpopulations of SECs, most frequently in *BRCA1/2* carriers, characterised by high *TP53* expression and markedly elevated histone levels. This phenotype is consistent with replication stress, cell-cycle arrest, and activation of innate immune signalling pathways. Protein-level validation in matched samples showed enrichment of cells with increased γ-H2AX expression and persistent 53BP1 foci in *BRCA1/2* carriers. Together our data supports the role of BRCA1/2 in maintaining genomic integrity and a *BRCA1/2* haploinsufficient phenotype characterised by increased replication stress in the fallopian tube epithelium. Our findings provide evidence for distinct immune responses in users of hormonal contraception and demonstrate early events in malignant transformation. They establish potential biomarkers in microscopically normal FT and a framework for measurement of cancer risk with the goal of enabling molecularly informed cancer interception in high-risk individuals.

## Introduction

Women carrying germline alterations in *BRCA1* or *BRCA2* face a markedly elevated lifetime risk of developing breast and ovarian cancers. Current prevention relies largely on risk-reducing mastectomy and salpingo-oophorectomy, which – despite their efficacy in reducing cancer risk – cause irreversible infertility, premature menopause, and substantial physical and psychological burden. Except for the protective effect of hormonal contraception (from retrospective studies), the development of alternative prevention strategies has been constrained by limited understanding of tumour initiation, particularly in high-grade serous ovarian carcinoma (HGSOC), whose origin in the fallopian tube (FT) epithelium has only been firmly established within the past two decades.

Previous efforts to investigate HGSOC initiation have focused on pre-neoplastic serous tubal intraepithelial carcinoma (STIC) lesions or bulk tissue comparisons between high-risk fallopian tubes and those from low-risk controls and have not provided an understanding of subtle gradients between cell states. Equally, single-cell profiling of fallopian tubes in healthy, low-risk individuals did not provide insight into early malignant transformation ^1–4^. Wider profiling of high-risk individuals across a spectrum of hormonal status is required, to fully understand the impact of combined genetic predisposition and hormonal regulation in HGSOC risk ^2,5–8^. Specifically, it is unclear how hormonal alteration by the menstrual cycle, menopause and from hormonal contraception alters the physiology and immunology of the fallopian tube epithelium. Finally, malignant transformation in the FT fimbria has been strongly associated with proximity to the ovary and ovulation-induced oxidative stress ^9^. However, understanding how individual cell types interact under different hormonal conditions and across different FT regions may reveal other factors that support the increased rate of malignant transformation in the fimbria.

Here, we analyse gene expression of 168,679 fallopian tube cells from the prospective TARGET-FAL01 study including 55 multiregional samples obtained from 34 women including 15 *BRCA1/2* mutation carriers to test associations between hormonal and risk status, immune microenvironment and evidence of cancer initiation signatures. Our findings provide a molecular framework for understanding the physiology of the fallopian tube in high-risk individuals, identify epithelial cell populations enriched in these individuals, and highlight potential targets for early intervention.

## Results

### Comprehensive atlas of normal fallopian tubes reveals significant heterogeneity and novel cell states

We collected tissue sections from normal fallopian tubes of 34 women aged 27–71 that underwent planned salpingectomy (Fig. 1a and Supplementary Table 1). From the 19 individuals undergoing risk-reducing salpingectomy, 15 had germline deleterious *BRCA1* or *BRCA2* mutations. The cohort included pre-, peri- and post-menopause (18, 3 and 13 subjects, respectively) in 30 parous and 3 nulliparous women. Menstrual cycle information was available for 18 pre- and peri-menopausal women. Only 5 women reported never having used hormonal contraception (HC) (Fig. 1b, c). To characterise the FT isthmus, ampulla and fimbria (proximal to distal from the uterus), we sampled tissue sections from more than one region from 14 subjects (Fig. 1d and Supplementary Table 2). We pooled multiregional samples across subjects into shared single-cell RNA-seq (scRNA-seq) libraries and computationally demultiplexed them by matching to subject genotypes obtained from each subject’s initial, unpooled sample (Supplementary Fig. 1a–f). We removed suspected doublets and discarded cells with less than 1500 unique molecular identifiers (UMIs) or with >20% expression of mitochondrial genes (Supplementary Fig. 1g–i). We also optimized sample handling to use single cell FT suspensions from frozen tissue (Supplementary Fig. 2a–b).

**Figure 1:**
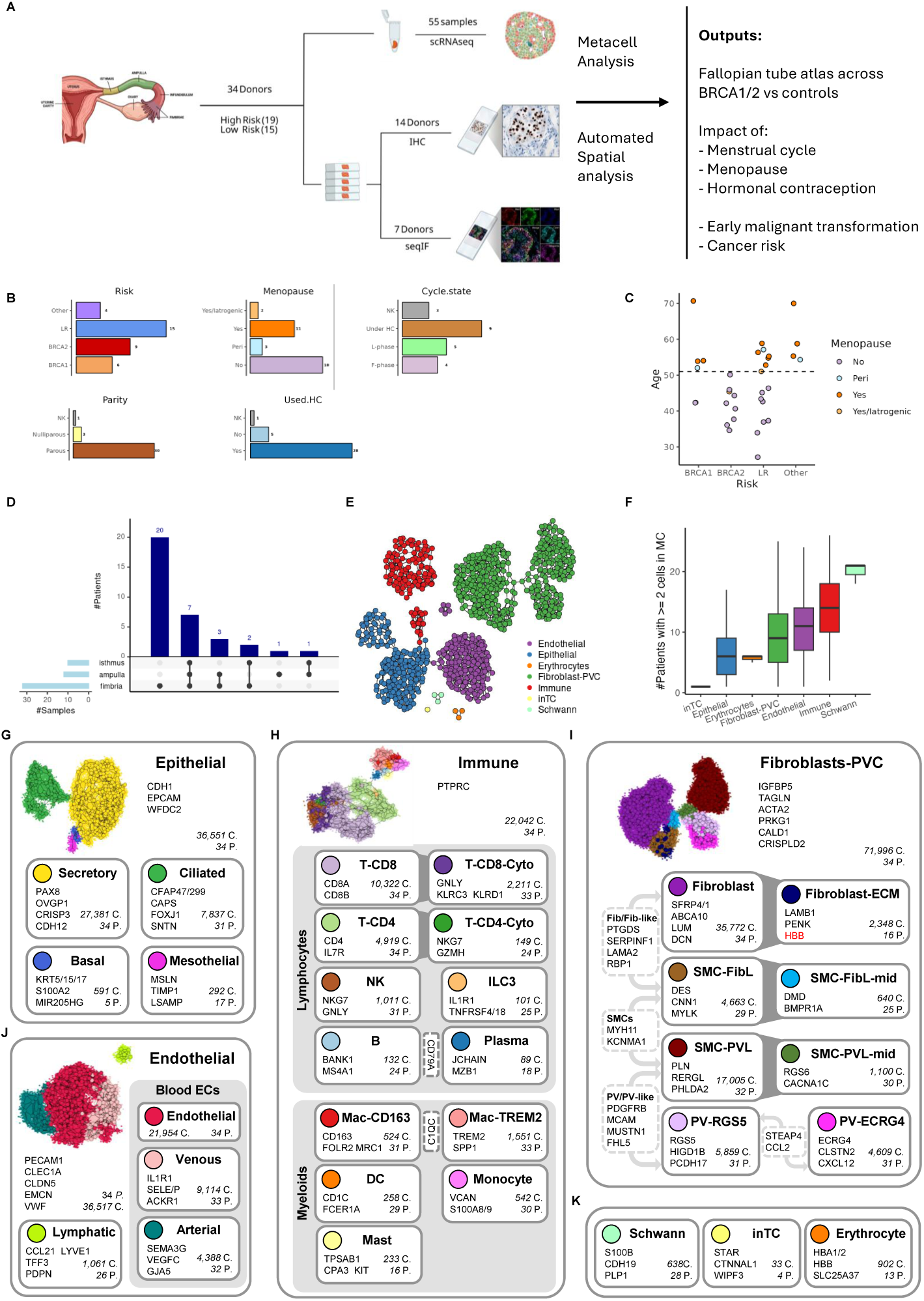
Metacell analysis of fallopian tube tissues. **a,** Study overview showing experimental design. scRNA, single cell RNAseq; p53 IHC, p53 immunohistochemistry on fallopian tube tissue sections; seqIF, sequential immunofluorescence on fallopian tube tissue sections. **b,** Barplots showing clinical features of study cohort. Numbers of subjects are indicated. **c,** Dot plot of age by risk category, coloured by menopausal status. Dashed line indicates average age at menopause in the UK. **d,** UpSet plot shows the number of subjects and regions of fallopian tube sampled. Numbers of subjects are indicated on the horizonal bar plot. **e,** 2D visualization map of 945 metacells grouping the 168,679 cells that passed QC. Metacells are annotated by high-level cell type. Transcriptionally similar metacells are connected by edges, otherwise position is arbitrary. **f,** Boxplot showing the number of subjects contributing cells to each metacell. **g,** 2D visualization map of the 197 epithelial metacells with their detailed cell-type annotation. Number of cells, subjects and marker genes are shown for each cell-type and for all epithelial cells. **h,** 2D visualization map of 127 immune metacells. Cell-types are grouped to the lymphoid and myeloid lineages, T-CD8-Cyto (cytotoxic CD8) and T-CD4-Cyto (cytotoxic CD4) are subsets of T-CD8 and T-CD4, respectively. Gene symbols common to cell-types are shown in dashed boxes. **i,** 2D visualization map of 416 fibroblast / peri-vascular metacells (PVC). **j,** 2D visualization map of 201 endothelial metacells. **k,** Cell type markers for Schwann, erythrocytes and interna theca (inTC) metacells. C, Number of cells; P number of subjects. Boxplots show median and the lower and upper hinges correspond to the first and third quartiles (the 25th and 75th percentiles). The upper whisker extends from the hinge to the largest value no further than 1.5 × IQR from the hinge (where IQR is the inter-quartile range, or distance between the first and third quartiles). The lower whisker extends from the hinge to the smallest value at most 1.5 × IQR of the hinge. Data beyond the end of the whiskers are plotted individually.

We reasoned that understanding diverse FT cellular states across physiological and pathological perturbations would require detection of subtle transcriptional gradients and small discrete transcriptional programmes. To obtain high granularity analyses we used *Metacell* ^10^ as this framework groups together highly similar cells that only differ as a result of sampling variance, thus providing homogeneous units of transcriptional states. Metacells form statistically robust building blocks for analysis ^11–14^ by preserving rare subpopulations rather than forcing them into larger, heterogeneous clusters.

We partitioned 168,679 cells derived from 55 samples into 945 metacells. We first annotated the metacells to broad cell-types by strongly expressed core transcriptional programs (Fig. 1e and Supplementary Fig. 2c). As expected, the main cell groups identified (Supplementary Table 7) were epithelial, immune, endothelial, and fibroblast / peri-vascular cells (PVC). We also observed other cell groups including erythrocytes from samples processed from fresh specimens and Schwann cells ^15^ which expressed *S100B*, *CDH19*, and *PLP1*. In subject FAL106, we identified a single metacell consisting of 33 cells expressing *STAR* and *CTNNAL1* consistent with ovarian interna theca cells ^16^ suggesting adherence of ovarian tissue to the fimbria (Fig. 1k and Supplementary Fig. 2a). As the number of subjects contributing cells to a metacell represent how common or specific is the cell state captured by that metacell, we first ordered metacells across the cohort by the number of contributing subjects. Schwann metacells captured the most invariant cell states, followed by immune metacells (Fig. 1f and Supplementary Fig. 2d). The stromal cell types showed increasing subject-specific phenotypes, from endothelial, through to fibroblast-perivascular cells (PVC). Epithelial metacells were the most subject specific, suggesting strong and dynamic relationships to physiological or pathological state.

We next refined cell-types and cell-states within each large cell group using more granular metacell annotation (Supplementary Table 7). Epithelial metacells (expressing *EPCAM*, *CDH1*, *WFDC2*) were further resolved into secretory (*PAX8*, *OVGP1*, *CRISP3*), ciliated (*CAPS*, *CFAP47*, *FOXJ1*), basal (*KRT5*, *KRT15*, *KRT17*) and mesothelial cells (*MSLN*, *TIMP1*) subsets (Fig. 1g, Supplementary Fig. 3a and Supplementary Fig. 4a). Lymphoid subtypes included CD4 and CD8 T cells, B cells, NK cells, plasma cells, whilst the myeloid lineage comprised monocytes, dendritic cells (DC) macrophages and mast cells (Fig. 1h and Supplementary Fig. 3b). We identified subtypes of cytotoxic CD4 (expressing *NKG7*, *GZMH*) and CD8 cells (expressing *GNLY*, *KLRC3*) (Supplementary Fig. 4g–h) and of type 3 innate lymphoid cells (expressing *IL1R1*, *TNFRSF4/18*) ^17^. Tumour-associated macrophages (expressing *TREM2*, *SPP1*) ^18^ were unexpectedly found in healthy FT tissue (Fig. 1h and Supplementary Fig. 4f) alongside the expected tissue resident macrophages (positive for *CD163*, *FOLR2*, *MRC1*). Fibroblast and PVC cells were very heterogeneous (Fig. 1i and Supplementary Fig. 5c). We observed a fibroblast sub-population expressing the extracellular matrix genes *LAMB1* and *PENK* (Fibroblast-ECM) in one subject, but this was excluded from further analysis. The perivascular (PV) metacells expressed *STEAP4* and *CCL2* and were further split into PV-RGS5⁺ and PV-ECRG4⁺ metacells characterised by *ECRG4*, *CLSTN2* and *CXCL12* expression (Supplementary Fig. 4e). All smooth muscle cells (SMCs) from this Fibroblast-PVC compartment expressed *MYH11* and KCNMA1 and resolved to fibroblast-like contractile (SMC-FibL, expressing *DES*, *CNN1* and *MYLK*) and PV-like vascular SMCs (SMC-PVL, expressing *PLN*, *RERGL* and *PHLDA2*) (Fig. 1i and Supplementary Fig. 4b), each with transitional states (SMC-Fib-mid; SMC-PVL-mid) bridging functional compartments (Supplementary Fig. 4c–d). Endothelial metacells comprised lymphatic (expressing *CCL21*, *TFF3*, *LYVE1*) and blood endothelial cells (*PECAM1*, *VWF*, *CLEC1A*), further stratified into arterial (*VEGFC*, *CXCL12*, *NEBL*) and venous (*SELE*, *ACKR1*) states (Fig. 1j and Supplementary Fig. 3d). Taken together, these data show that using a metacell approach we detect unexpected subject heterogeneity in normal fallopian tube epithelia and its microenvironment, as well as confirming previously observed cell populations ^2^ (Supplementary Fig. 6). These analyses provided the critical ground state to explore more granular annotation of cell populations and states from physiological and hormonal perturbations.

### Menopausal effects on fallopian tube serous epithelial cells

We took two complementary approaches to explore how hormonal status and exposure affect fallopian tube cell composition and gene expression. First, differential abundance analysis identified shifts in cell-type frequencies across patient groups. Second, composition-balanced differential expression analysis revealed within-cell-type transcriptional changes. We controlled for confounding variables such as risk status and anatomical region (Supplementary Table 8) and used leave-one-out analyses to mitigate differences arising from a single patient (see Methods).

Consistent with earlier reports, ^2,19^ menopause was associated with depletion of ciliated epithelial cells (CECs) (Fig. 2a). We observed other reported compositional changes^2^ including depletion of secretory epithelial cells (SECs) and increase of PVs and SMC-PVL, but these changes were not statistically significant in our data (Supplementary Fig. 7a). Composition-balanced differential expression analysis of post-vs pre-menopausal samples in subjects not using hormonal therapy identified major changes in SECs (Supplementary Fig. 8a–b), with lower post-menopausal expression of key SEC markers (*OVGP1*, *WFDC2*) and higher expression of some signalling genes (*AKAP12*, *SLIT3*, *DOK5*) (Fig. 2b and Supplementary Table 9). We also observed shared phenotypic changes across many cell types, predominantly increased stress response genes (*FOSB*, *ATF3*) and specific oxidative stress response genes (*MT1A/X*, *CEBPD*), which have been reported in menopause ^2,3^. Interestingly, the expression of *MTRNR2L8*, which is also an oxidative stress response gene, was lower after menopause across most cell-types, as previously reported ^2^. We did not observe the previously reported decrease in expression of ribosomal protein genes upon menopause in low-risk subjects. However, in *BRCA1/2* subjects, we saw a strong increase in ribosomal protein gene expression across multiple cell-types, most pronounced in SECs (Supplementary Fig. 8c–e). These data show that menopause predominantly impacts the phenotypic profile of SECs, the precursor population of HGSOC.

**Figure 2:**
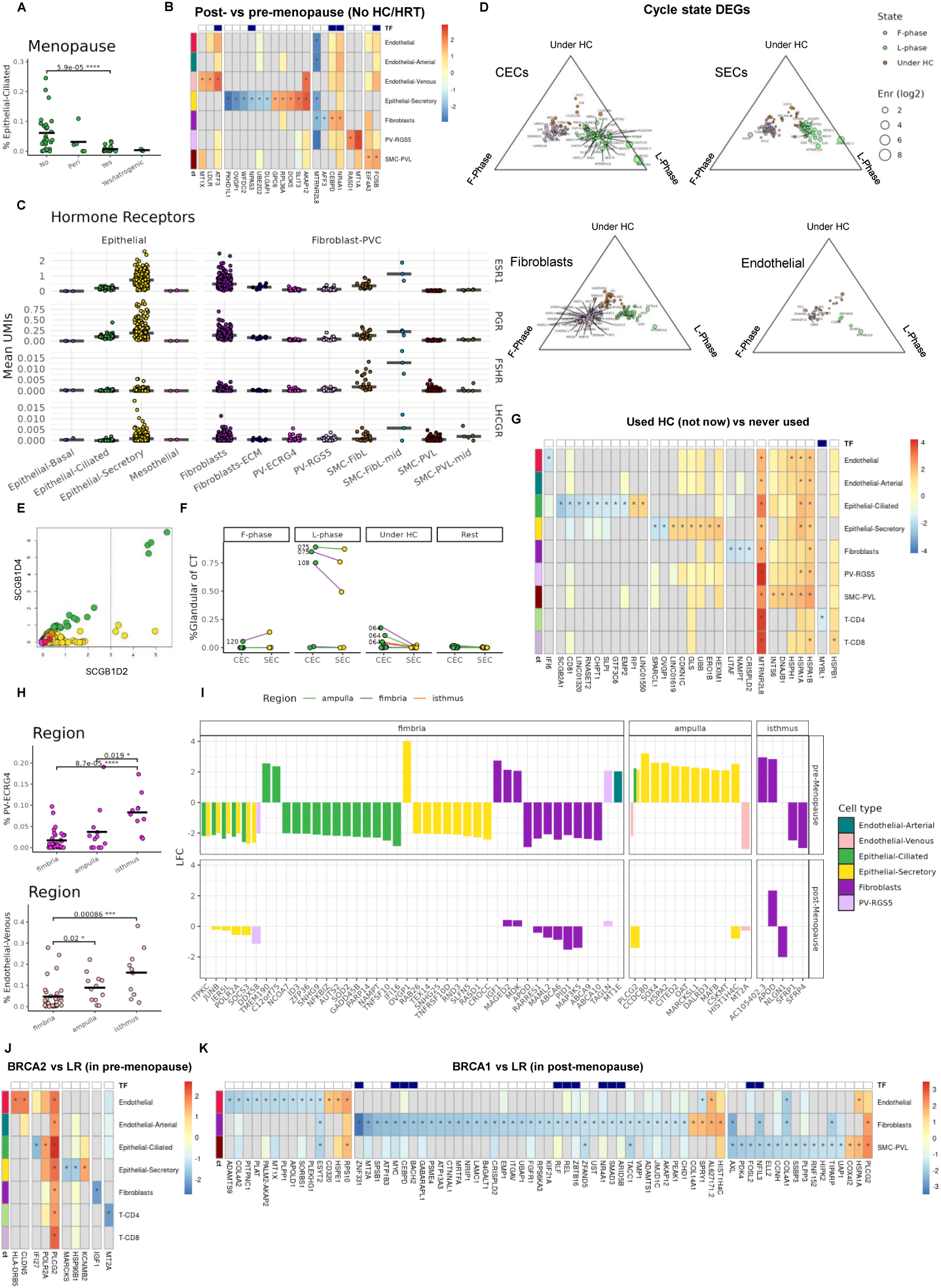
Effects of menopause, menstrual cycle and hormonal therapy on the fallopian tube. **a,** Dot plot showing fraction of ciliated epithelial cells (CEC) in samples by their menopause status, p-value by Mann-Whitney test. Horizontal bars indicate median proportion. **b,** Heatmap showing controlled differentially expressed genes (DEG) from post-vs pre-menopausal samples from subjects not using hormonal therapy (HC and HRT). Heatmap indicates log-fold change (LFC) and asterisks indicate absolute LFC ≥ 1. TF, transcription factor; ct, metacell cell type. **c,** Dot plots showing expression (count per 1000 UMIs) of hormone receptor genes across the epithelial and fibroblast/PVC metacells. **d,** Ternary plots showing effects of luteal and follicular phase of the menstrual cycle or hormonal therapy on genes that were enriched in the controlled DEG analysis. Gene symbols are shown for absolute LFC ≥ 1.75 or if gene count was ≤ 12. L-phase, luteal phase; F-phase, follicular phase. **e,** Metacell gene enrichment (log2) for the *SCGB1D2* and *SCGB1D4* secretoglobins. Metacells are coloured by their detailed annotation show in Fig. 1g–k. Vertical dashed line indicates cutoff defining glandular-CEC (gCEC) and glandular-SEC (gSEC). **f,** Effects of luteal and follicular phase of the menstrual cycle or hormonal therapy of the fraction of gCEC and gSECs. Rest indicates pooled non-pre-menopausal samples. Lines connect cell-types from the same subject. Labels show subject numbers. **g,** Heatmap showing controlled DEG between samples from subjects who previously used hormonal contraception (HC) and subjects with no HC exposure. Heatmap indicates log-fold change (LFC) and asterisks indicate absolute LFC ≥ 1. Ct, metacell cell type **h,** Fraction of PVC-ECRG4 and venous endothelial metacells in fallopian tube regions. **i,** Bar plots showing DEGs across FT regions in pre- and post-menopausal subjects. Genes are filtered within each metacell type by LFC ≥ 2. **j,** Heatmap comparing samples from pre-menopausal *BRCA2* vs low-risk (LR) subjects, showing genes with absolute LFC ≥ 1.5. Ct, metacell cell type. **k,** Heatmap comparing samples from post-menopausal *BRCA1* vs LR subjects.

### Menstrual cycle and hormonal regulation of the fallopian tube

Unlike the endometrium and ovary, hormonal regulation of individual FT cell types has not been extensively studied. Epidemiological data show that the number of lifetime ovulations is directly associated with HGSOC risk, with both oral contraception and pregnancy being protective ^20^. Since the mechanisms for this protection remain unclear, we investigated the role of hormone therapy in regulating proliferative FT epithelial cells or cells from the stromal compartment.

During the follicular phase, follicle-stimulating hormone (FSH) promotes follicular development and drives a progressive rise in oestrogen levels. Towards the end of the follicular phase, peak oestrogen levels, together with FSH, trigger a surge in luteinising hormone (LH), resulting in ovulation. The luteal phase is characterised by high progesterone levels, which, together with oestrogen, decline if pregnancy does not occur. Metacell analysis confirmed that SECs and fibroblasts are the primary cell types expressing oestrogen (*ESR1*), progesterone (*PGR*) and luteinising hormone (*LHCGR*) receptors, whereas elevated *FSHR* (FSH receptor) expression was restricted to the SMC-Fib and SMC-Fib-mid populations (Fig. 2c and Supplementary Fig. 9a–b). Interestingly, SMC-FibL-mid were more abundant in luteal-phase, never occur in follicular-phase (Supplementary Fig. 7b) and exhibit increased abundance along the FT, peaking at the isthmus (Supplementary Fig. 10a). In luteal-phase, samples with high numbers of glandular CECs (see below) also had a high abundance of SMC-FibL-mid cells (Supplementary Fig. 9c), suggesting that this represents a transient cell state occurring after FSH surge. Only *PGR* and *ESR1*, but not *LHCGR* or *FSHR,* showed differential receptor expression by cycle state or menopause (Supplementary Fig. 9d–e). *PGR* expression was decreased in SECs upon menopause and in pre-menopausal women using hormonal contraception. This may be the result of lower circulating estradiol upon menopause or through progestins-induced suppression in the hypothalamic-ovarian axis. Conversely, we identified increased *PGR* and *ESR1* in follicular-phase fibroblasts which may represent result from follicular unopposed estrogen driving classical positive-feedback gene expression before ovulation.

In pre-menopausal women – the stage of menstrual cycle, follicular-phase or luteal-phase, and concurrent use of hormonal contraception impacted a wider number of cell types including CECs, fibroblasts and to a lesser extent endothelial cells (Fig. 2d and Supplementary Fig. 11a–c). Not surprisingly, we also observed co-ordinated regulation during the menstrual cycle across CECs, SECs and fibroblasts (Supplementary Fig. 11d). Follicular-phase samples were enriched for the *CRY1* circadian gene (in SECs and fibroblasts), *CRIM1* and *SOX4* (in SECs and CECs ^21^), and early response genes *EGR1*, *JUN* (simultaneously in SECs, CECs and fibroblasts). Luteal-phase samples were enriched for the epithelial markers *WFDC2* and *SLPI* in CECs and SECs and *GSN* in all cell types. *MTRNR2L8* and the long non-coding RNA *AC105402.3* gene were also primarily enriched in luteal SECs and CECs in some subjects (Supplementary Fig. 11e) ^22^. Fibroblasts displayed distinct transcriptional profiles across the menstrual phases (notably *MGP*, *GSN* and *TAGLN* enrichment in the luteal-phase and *SFRP4* in the follicular-phase; Fig. 2d). These findings demonstrate that, prior to menopause, hormonal regulation during the menstrual cycle and changes associated with hormonal contraception have broad cross-cellular phenotypic effects, influencing pathways involved in DNA damage responses, cell-cycle regulation, and cellular differentiation.

The glandular secretoglobulin markers *SCGB1D2* and *SCGB1D4* have been previously shown to be enriched in the luteal endometrium ^23^, and we observed strong enrichment in luteal-phase CECs (Fig. 2d). We used *SCGB1D2* expression to define glandular-CECs (gCEC) and SECs (gSEC) (Fig. 2e). Two subjects, both in luteal phase, showed a strong shift to glandular phenotype in most of their SECs and CECs (Fig. 2f). These cell populations expressed *RNASE1*, suggesting a defensive role during luteal-phase against extra-cellular RNA in the fallopian tube (Supplementary Fig. 11f–g).

### Use of hormonal contraception has persistent effects on the fallopian tube epithelia and alters MHC expression

Concurrent use of hormonal contraception was associated with increased expression of stress response and cycle arrest related genes (Supplementary Fig. 11a–b). Equally, previous use of hormonal contraception (HC) had a detectable expression phenotype, even after controlling for FT region, menopausal and risk status (Fig. 2g). 18 women with prior HC exposure, when compared to 5 women who never used HC, showed reduced expression of lineage-defining markers (*OVGP1* in SECs, *SLPI* and *SCGB2A1* in CECs and *CRISPLD2* in fibroblasts), and a cross cell-type increase in stress response genes (*DNAJB1*, *INTS6* and heat shock protein genes). Moreover, CD163-positive tissue resident macrophages were more frequent in those who had never used HC (Supplementary Fig. 9f), demonstrating potential lasting effects of hormonal contraception usage in both FT epithelium and immune surveillance.

We next focused on how hormonal regulation impact the innate and adaptive immune systems ^24^. Surprisingly, all venous endothelial cells and a small subset of SECs expressed MHC class II genes at comparable levels to antigen presenting immune cells (Supplementary Fig. 12a–b). Venous endothelial cells trigger immune responses during inflammation ^25^, and MHC class II-positive SECs subsets have previously been reported in the FT ^2,26,27^. In our dataset, MHC class II gene expression in SECs and CECs is lowest during follicular-phase ^27^, then increases in luteal-phase and reaches its highest levels during exposure to hormonal contraception. (Supplementary Fig. 12c–h). Notably, the SECs with the highest MHC-II expression were from women using progesterone-based HC, either via injection or an intrauterine device, with a similar but weaker pattern seen in CECs. Gene set enrichment analysis of SEC metacells expressing MHC-II showed broad activation of immune pathways and depletion of cell cycle genes (Supplementary Fig. 12i). Notably, we found coordinated depletion of MHC-II genes across SECs, CECs and endothelial cells (ECs) in the pre-menopausal fimbria in low-risk individuals but strong enrichment in pre-menopausal *BRCA2* carriers. Critically, previous use of hormonal contraception was associated with enrichment of MHC-II expression in SECs and depletion in ECs (Supplementary Fig. 12j).

### Regional specification of the tubal epithelium

The anatomical regions of the FT (fimbria, ampulla, and isthmus) have distinct physiological functions, but detailed molecular characterization has not been extensively performed. We did not find subject-specific cross-region differences in cell-type abundance, as regional samples from the same subject resembled each other as much as those from different people (Supplementary Fig. 10b–c). The isthmus had the least inter-subject variation in cell-type composition. Consistent with previous reports we did not observe region-exclusive cell-types ^3,4^. We observed a gradual increase from the fimbria to the isthmus for both *PV-ECRG4* and venous ECs, likely related to an increase in vessel size (Fig. 2h). Controlled cross-region differential expression revealed regional cell-type specialisation. In the fimbria, CECs had increased expression of the cilia machinery genes *TMEM190*, *C12orf75* and SECs had higher expression of *OVGP1*, enhancing support and transport for oocyte movement through the FT ^28^. Fimbria fibroblasts had increased expression of growth and repair genes (*IGF1*, *MDK*, *MAGED2*) (Supplementary Fig. 10d–e). Ampullary SECs expressed genes associated with metabolism and nutrient production (*OAT*, *CSKMT*, *DALRD3*), stress defence genes (*HSPA2*, *CITED2*), fluid secretion (*MARCKSL1*, *CCDC80*), and epithelial proliferation genes (SOX4, MAFB, HIST1H4C), consistent with a role for maintaining the oocyte for fertilization. Both ampullary SECs and CECs had significant upregulation of *PLCG*2 (see next section) ^29^. After menopause, regional differences were diminished but consistent with pre-menopausal specification patterns (Supplementary Fig. 8a–b).

### Carriers of BRCA1/2 germline alterations express putative tumorigenic genes across many cell-types

As previously shown, we did not find significant differences in cell-type composition between *BRCA1/2* mutation carriers and non-carriers. To investigate transcriptional differences between these groups, we compared *BRCA1/BRCA2* FTs with controls, balancing for menopausal status, hormonal exposure, sample region and for cycle state in pre-menopausal subjects (Fig. 2j–k and Supplementary Fig. 13a–b). Across multiple epithelial and stromal cell types, we observed strong enrichment of *PLCG2, MTRNR2L12* (long-coding RNA; pseudogene) and *HIST1H4*C in both *BRCA1* and *BRCA2* carriers. Notably, *PLCG2,* which has calcium-mediated roles in proliferation and cell signalling through exo- and endocytosis in both epithelial and immune cells, and *MTRNR2L12* were recently identified as marking cellular subpopulations associated with poor prognosis and metastasis in small cell lung cancer ^29,30^. In our cohort, positive expression of these genes was correlated with active use of hormonal contraceptive (Supplementary Fig. 11b and Supplementary Fig. 13c–d) and was particularly enriched in pre-menopausal ampullar SECs and CECs (Fig. 2I). Interestingly, both gCECs and gSECs shared extreme depletion of *PLCG2* and *MTRNR2L12* in comparison to their non-glandular counterparts (Supplementary Fig. 11f–g). The broad, cross-cell-type enrichment of *PLCG2* and *MTRNR2L12* in *BRCA1/2* carriers highlights the functional impacts of germline mutation on FT and suggests that transcriptional alterations predate overt neoplasia, marking an early state of vulnerability.

### A sub-population of secretory epithelial cells enriched in *TP53* expression may reflect a hallmark of *BRCA1/2* haploinsufficiency

We hypothesised that aneuploidy may arise sporadically in healthy fallopian tube (FT) epithelium of BRCA1/2 pathogenic variant carriers at higher rate than in low-risk individuals, and that such events would be associated with activation of repair mechanisms, cell-cycle checkpoints, and the p53 pathway. The p53 protein, and its coding gene *TP53*, are of special interest, as “p53 signatures” (clusters of 10–15 SECs with mutant p53 accumulation ^31^) are precursors of serous tubal intra-epithelial carcinoma (STIC), invasive high-grade serous ovarian carcinoma (HGSOC) and show significant chromosomal instability with numerous chromosomal gains and losses ^32^. We therefore used our metacell analysis to test for surrogate measures of chromosomal instability and identified a rare subset of SEC metacells with elevated *TP53* expression and concomitant activation of both G1/S and/or G2/M transcriptional programs (Fig. 3a–b). Since mRNA expression of *TP53* is known to be high in both G1/S and G2/M checkpoints, we used both signatures ^33^ to define cutoffs on metacells (Supplementary Fig. 14a–c), grouping them to G1/S ^+^, G2/M ^+^, negative for both and positive for both (G1/S ^+^–G2/M ^+^). These double-positive G1/S ^+^–G2/M ^+^ metacells are rare, appearing in frequency higher than 2% in only 8 of our samples, from 7 subjects, 5 *BRCA2* and one *BRCA1* carrier (Supplementary Fig. 15), which may explain their absence from previous FT scRNA-seq datasets that have low numbers of high-risk patients ^1,2,4,34^. Their frequency and transcriptional profile resemble proliferative states in HGSOC ^35–38^ (Supplementary Fig. 14d–f). All the G1/S ^+^–G2/M ^+^ metacells were SECs (Fig. 3b and Supplementary Fig. 14g), expressing markers from different cell-cycle phases consistent with hyper or abnormal proliferation: G1/S markers (*CCNE1/2*, *CDC25B*), G2/M (*CCNB1*, *CDC25C*, *NUSAP1*) and M-phase (*CENPE*) (Supplementary Fig. 16).

**Figure 3:**
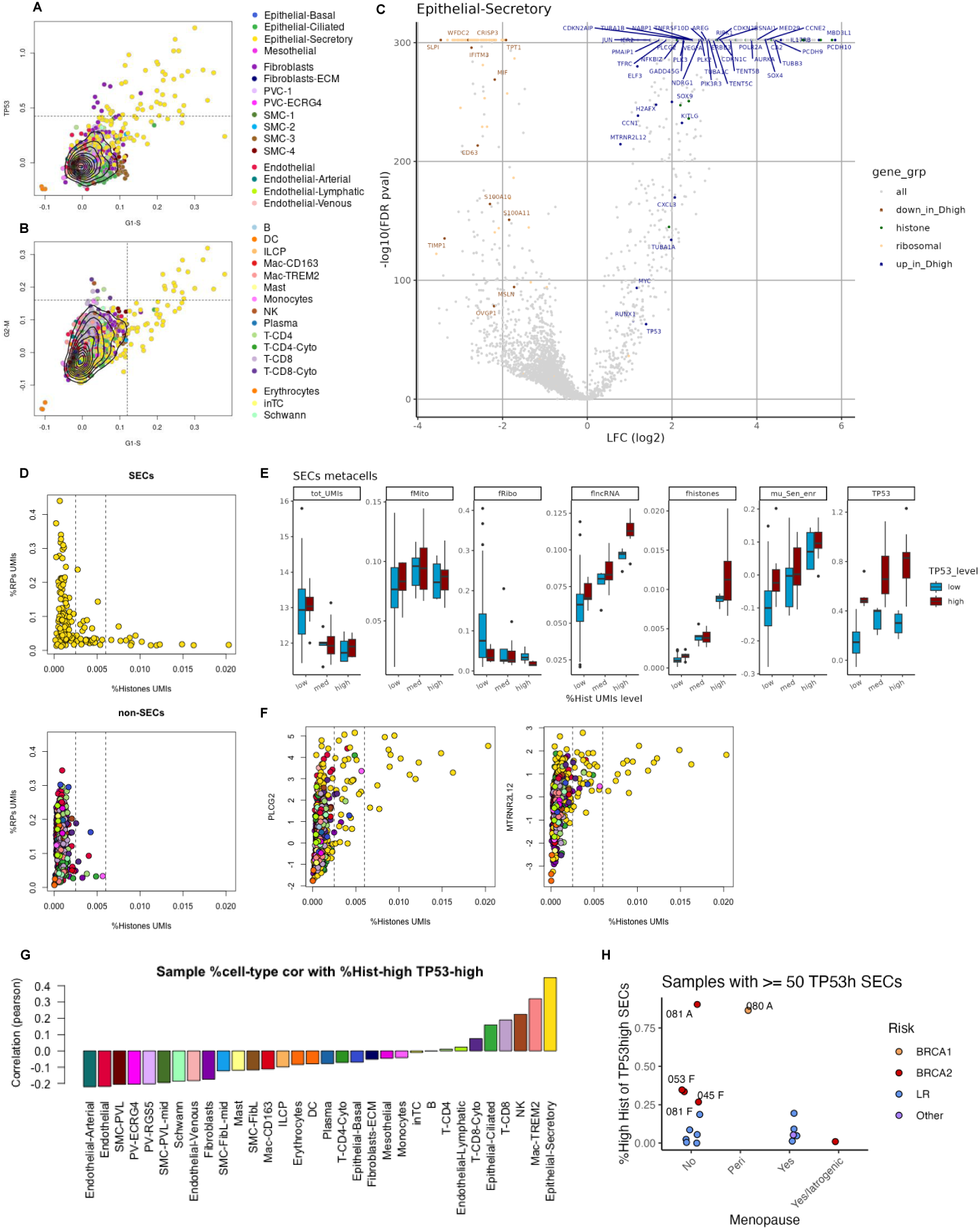
A sub-population of secretory epithelial cells are enriched in *TP53* expression. **a,** *TP53* gene expression per metacell versus mean G1–S gene expression. Dashed line indicates *TP53*-high cut point for metacells. **b,** Mean G1–S expression per metacell versus mean G2–M gene expression. Dashed lines indicate cut points for G1–S and G2–M positive metacells. **c,** Volcano plot of DEGs comparing G1–S ^+^/G2–M ^+^ SECs vs G1–S ^−^/G2–M ^−^ SECs. Legend indicates gene function. D-high indicates double-positive G1–S ^+^/G2–M ^+^ SECs. **d,** Mean fraction of UMIs from ribosomal protein genes and histone coding genes in SEC and non-SEC meta cells. Dashed lines indicate boundaries for low–mid and mid–high histone gene expression. **e,** Boxplots showing characteristics of SEC metacells within low, medium and high histone gene and *TP53* expression groups. Panel labels indicate tot_UMIs (log2 mean total UMIs), mean %UMIs from mitochondrial genes (fMito), ribosomal genes (fRibo), lncRNA genes (flncRNA), histone genes (fHistones), mean senescence gene enrichment ^42^ and *TP53* enrichment. **f,** *PLCG2* and *MTRNR2L12* log2 enrichment versus fraction of UMIs from histone genes per metacell. Dashed lines indicate boundaries for low–mid and mid–high histone gene expression. **g,** Barplots showing Pearson correlation coefficients between cell-type frequency and fraction of Histone ^high^-*TP53* ^high^ cells. **h,** Dot plot showing fraction of histone ^high^-*TP53* ^high^ SECs compared to *TP53* ^high^ SECs grouped by menopausal status and coloured by risk. Pre-menopausal BRCA2 samples are perfectly separated from pre-menopausal LR ones (p.value 0.0095 by two-sided Mann-Whitney test). Data represents samples ≥ 50 *TP53* ^high^ SECs. Subject number and FT region are indicated for samples with highest histone fraction. F, fimbria; A, ampulla.

Differential expression analysis (Fig. 3c and Supplementary Fig. 17a) showed that these G1/S ^+^–G2/M ^+^ SECs were enriched in stress response genes (*JUN*, *FOSB*, *EGR1*), genes associated with cell-cycle arrest, cell death and senescence (*CDKN1C*, *CDKN2B*, *RIPK1*, *TENT5C*, *PMAIP1*, *H2AFX*), and transcription machinery genes (*POLR2A*, *MED29*). These SECs exhibit transcriptional de-differentiation, with reduction of epithelial genes (*CRISP3*, *WFDC2*, *SLPI*, *MSLN*, *OVGP1*) and increase in genes markers of other lineages, such as mesenchymal (*SNAI1*, *VIM*, *FN1*) and neuronal (*PCDH10*, *TUBB3*, *NRIP3*, *TFRC*), in line with what has previously been seen in the context of abrogated *TP53* ^39^. These SECs also showed *MYC* and *RUNX1* enrichment indicating abnormal proliferation. Strikingly, they exhibited a pronounced enrichment of canonical histone coding genes and depletion of ribosomal protein genes (Fig. 3d), with a subset of “histone-high” SECS having the highest expression of *TP53,* aligning precisely with G1/S ^+^–G2/M ^+^ SECs (Supplementary Fig. 17b–e). Histone production is a rate limiting step for S-phase progression and histone hypertranscription, characterised by poly-adenylation of histone mRNAs, is influenced by replication stress and can drive hyperproliferation and aneuploidy in cancer ^40,41^. Further detailed analyses of SEC metacells with the highest *TP53* and histone expression showed strong associations with transcriptional de-differentiation, stress, cell-death and senescence genes (Supplementary Fig. 18), with lower mRNA counts, higher levels of mitochondrial and lncRNA (Fig. 3e). These data suggest that TP53/histone ^high^ SEC metacells include cells under significant replication stress, which can progress through abrogated G2/M checkpoints or arrest at either G1/S or G2/M checkpoint ^42^. In addition, TP53/histone ^high^ SEC metacells had higher expression for the BRCA-associated *PLCG2* and *MTRNR2L12* genes (Fig. 3c, f). Estimates of SNV clones (see below) were not increased in TP53/histone ^high^ SECs, which suggest they are not actively proliferating (Supplementary Fig. 17f).

TP53/histone ^high^ SECs were also the main cells expressing *GSDMA* (encoding Gasdermin A), suggesting activation of the transcriptional programs that drive pyroptotic cell death ^43^, an inflammatory process which could lead to recruitment of immune cells. In our cohort, the TP53/histone ^high^ SECs were also the most enriched for Mac-TREM2, NK cells and T-CD8 cells, supporting simultaneous recruitment of the innate immune system (Fig. 3g), as a homeostatic mechanism to eliminate cells undergoing replication stress. Furthermore, we found that only subset of TP53 ^high^ metacells had high cGAS expression (Supplementary Fig. 17g), suggesting variation in ability to initiate the signalling responsible for recruitment of innate immune system. This may suggest that some *TP53* ^high^ cells with the highest histone expression may already have *TP53* mutations and impaired ability to initiate these recruitment processes.

Haploinsufficiency of *BRCA1/2*, associated with chromosomal mis-segregation in the mammary epithelium ^44^, leads to an expansion of proliferative luminal cells harbouring chromosomal gains and losses ^45^. Pre-menopausal *BRCA1/2* carriers had the highest proportion of elevated *TP53*/histone ^high^ SECs (across *TP53* ^high^ SECs) in the fallopian tube (Fig. 3h) further supporting a cross-tissue *BRCA1/2*-associated haploinsufficiency phenotype and linking this aberrant cell population to risk of early malignant transformation.

### SECs show the highest levels of genomic instability from scRNA-seq analysis of single nucleotide variants and somatic copy number alterations

Normal tissues accumulate mutations with age, ^46^ but whether these changes occur in the FTs, cluster in specific cell types or link to clinical features is unclear. Following observations of chromosomal instability in normal breast tissue from high-risk individuals and our findings of sparse TP53/histone ^high^ SEC, we analysed the FT scRNA-seq to ask if there were detectable single-nucleotide variants (SNVs) or copy-number alterations (CNAs). As mutation detection from 3′-UTR biased scRNA-seq data is challenging owing to sparse and partial coverage we used SComatic ^47^ which filters false positives using normal tissue references and identifies lineage-specific SNVs (Supplementary Table 10). Across 27 subjects, we identified 622 SNVs (median 15 per subject, range 1–71) with no observable difference between *BRCA1/2* and low-risk subjects (Fig. 4a). These SNVs appeared in 1,631 cells, mostly SECs (Fig. 4b and Supplementary Fig. 19a–b). A single SNV was detected in almost all cells and 61 cells harboured ≥2 SNVs. SNVs were relatively depleted in the fimbria (Fig. 4c and Supplementary Fig. 19c). As SNVs are accumulated during replication cycles, this region-specific depletion might reflect high cell-turnover in the fimbria ^48^. SNV frequency was not correlated with subject age (Supplementary Fig. 19d). The largest SNV clones (with ≥5 cells sharing the same SNVs) were comprised of epithelial cells (Fig. 4d), particularly in *BRCA2* subjects (Supplementary Fig. 19e) and were often confined to a single sampled region (Fig. 4e). When assessing major cell compartments (epithelial, immune, fibroblast-PVC and endothelial) separately and correcting for cell number across cell types, SNVs in the fibroblast-PVC compartment tended to occur more frequently than expected in SMC-FibL and Mac-TREM2. Equally, epithelial SNVs tended to co-occur more frequently within each cell type with less SNVs being shared between SECs and CEC than expected by chance (Supplementary Fig. 19f). These findings could be related to the increased proliferative activity of SEC clones (Fig. 4f) and limited changes in cell state in SMC-Fibl and Mac-TREM2 but may also represent limitations of SNV calling from sparse scRNA-seq data.

**Figure 4:**
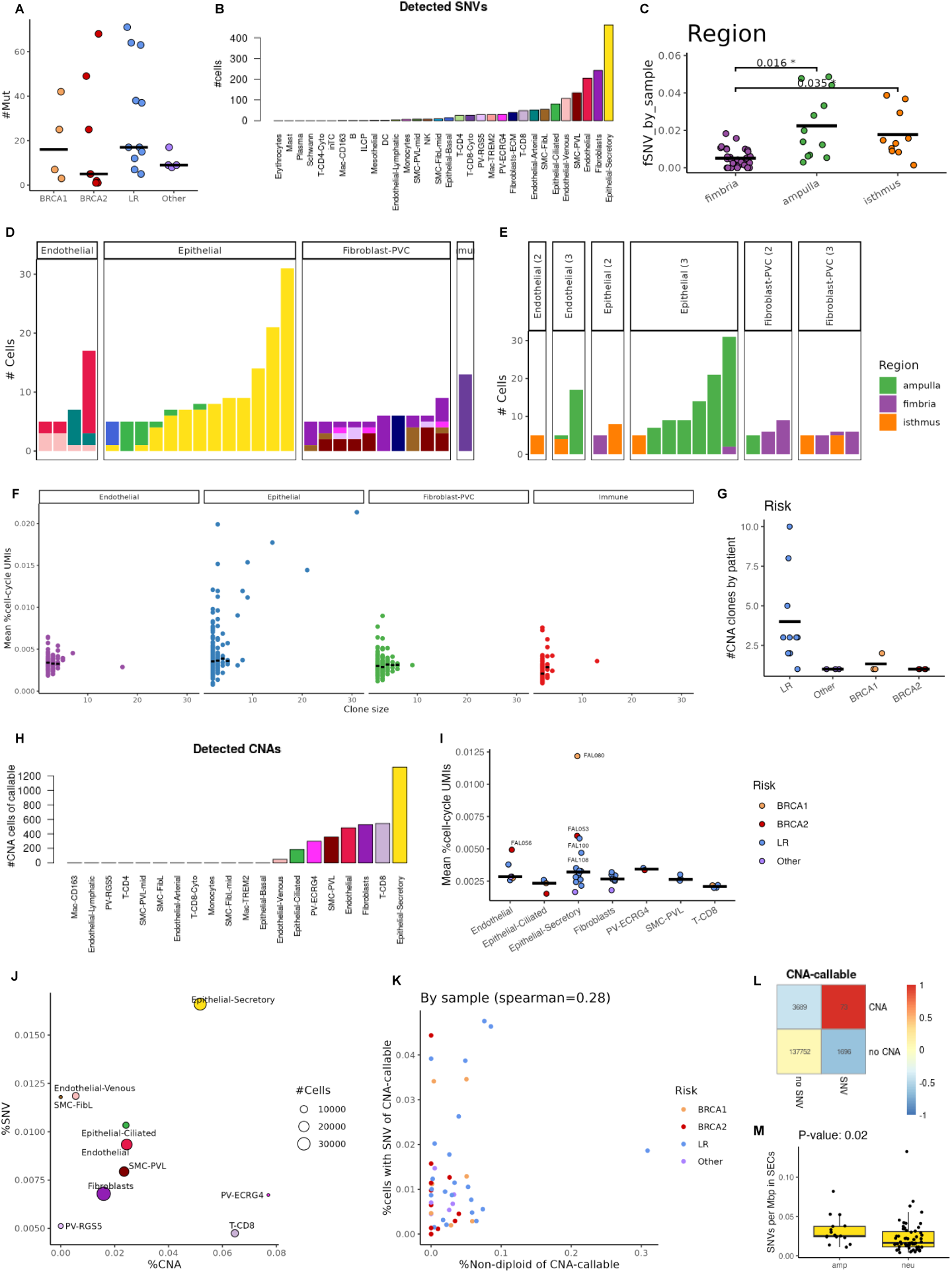
Analysis of single nucleotide variants and somatic copy number alterations in fallopian tube metacells. **a,** Dot plots showing number of distinct SNVs detected per subject grouping subjects by risk. **b,** Bar plot showing total number of cells with a detected SNVs by metacell. **c,** Dot plots showing fraction of cells with a detected SNV by sample grouped by FT region. Statistical significance was defined adjusting for multiple comparisons using the false discovery rate (FDR) and the Mann-Whitney test. **d,** Bar plots showing cell-type composition of SNV clones with ≥ 5 cells. Metacell type is indicated as in **b** and grouped by high-level cell-type and ordered by clone size. **e,** Dot plots showing regional FT composition of SNV clones with ≥ 5 cells from multi-region subjects. Panels indicate high-level cell-type and number of regions profiled per subject. **f,** Scatter plots showing fraction of UMIs from cell-cycle genes by clone size per SNV clone. Panels indicate high-level cell-type and median value for clone sizes is shown by black bars. **g,** Dot plots showing number of SCNA clones detected per subject grouped by risk. **h,** Bar plot showing total number of cells with a detected SCNA by cell-type. **i,** Dot plots showing fraction of UMIs from cell-cycle genes in SCNA clones grouped by cell-type and risk. Subject numbers are shown for highest clones. Horizontal bars indicate median values. **j,** Fraction of SCNAs and SNVs per cell-type. Number of cells is indicated as size of point. **k,** Comparison of the fraction of cells with any SCNA (non-diploid) and any SNV from cells that are SCNA callable (see Methods) by sample and coloured by risk. Spearman correlation is indicated. **l,** Contingency table for cells with SCNA and SNV from SCNA-callable cells. Colour indicates the z-score over expected numbers by chance. **m,** Boxplots showing SNV density per Mbp in SECs stratified by amplified or neutral SCNA state (*P*-value by Mann-Whitney test).

We looked for evidence of chromosomal instability in FT by using numbat ^49^, which integrates allelic SNPs to refine somatic copy number alteration (SCNA) detection from scRNA-seq (Supplementary Fig. 20a). We examined for evidence of clones in each subject and cell-type, using matched cell-type cells from all subjects as a normal reference. In 20 subjects, we identified 51 clones (1–10, median 1.5 per subject), with low-risk individuals carrying more clones, albeit with smaller clone sizes (Fig. 4g, Supplementary Table 11 and Supplementary Fig. 20b). SEC harboured the majority of the SCNA clones (Fig. 4h–i), without evidence of proliferative advantage (Supplementary Fig. 20c–e), suggesting they are stable. At multiple levels, including cell-type, sample, cell and genomic locus, we observed co-occurrence of SNVs and SCNAs (Fig. 4j–m).

Interestingly, the subjects who had the larger number of epithelial SNV clones (FAL081 and FAL053; Supplementary Fig. 19e) and highest cell-cycle activity in SCNA clones (FAL080 and FAL053; Fig. 4i), also had the highest proportion of elevated TP53/histone ^high^ SECs across *TP53* ^high^ SECs (Fig. 3h).

To further characterize these findings, we profiled 971 unselected cells from one high-risk and one low-risk normal FT using DLP+ single-cell DNA-seq (Supplementary Fig. 21) ^50^. Although these samples were not enriched for epithelial cells, we observed SCNA in 84 of the cells (8.7%). We observed recurrent loss of chromosome X which was also found across breast cell types and preferentially affected the inactive X chromosome, likely representing a selectively neutral event. We did not detect any large clonal SCNA expansions. These data are consistent with findings in normal breast epithelium, where rare isolated aneuploid cells were observed and the largest clone, harbouring a 1q gain, accounted for only 1.5% of luminal progenitor cells while being virtually absent from all other epithelial cell types.

### BRCA-associated FT fimbria show cellular changes driven by replication stress

Based on the TP53/histone ^high^ SEC metacell analyses, we next investigated spatial patterns of p53 accumulation through immunohistochemistry (IHC) in 259 FT sections from our subjects, representing all FT regions. Cells were classified as epithelial, stromal, or red blood cells after nuclei segmentation. Intensity of p53 expression was categorized as negative, low (1+), medium (2+), or high (3+) (Supplementary Table 1 and Fig. 5a). Tissues were manually annotated as fimbria or non-fimbria (Supplementary Fig. 22a–b), yielding 79 fimbria, 165 non-fimbria and 15 fragments from unknown regions (Fig. 5b), with *BRCA1/2* subjects having additional sampling as part of the SEE-FIM protocol (Fig. 5c and Supplementary Fig. 22c).

**Figure 5:**
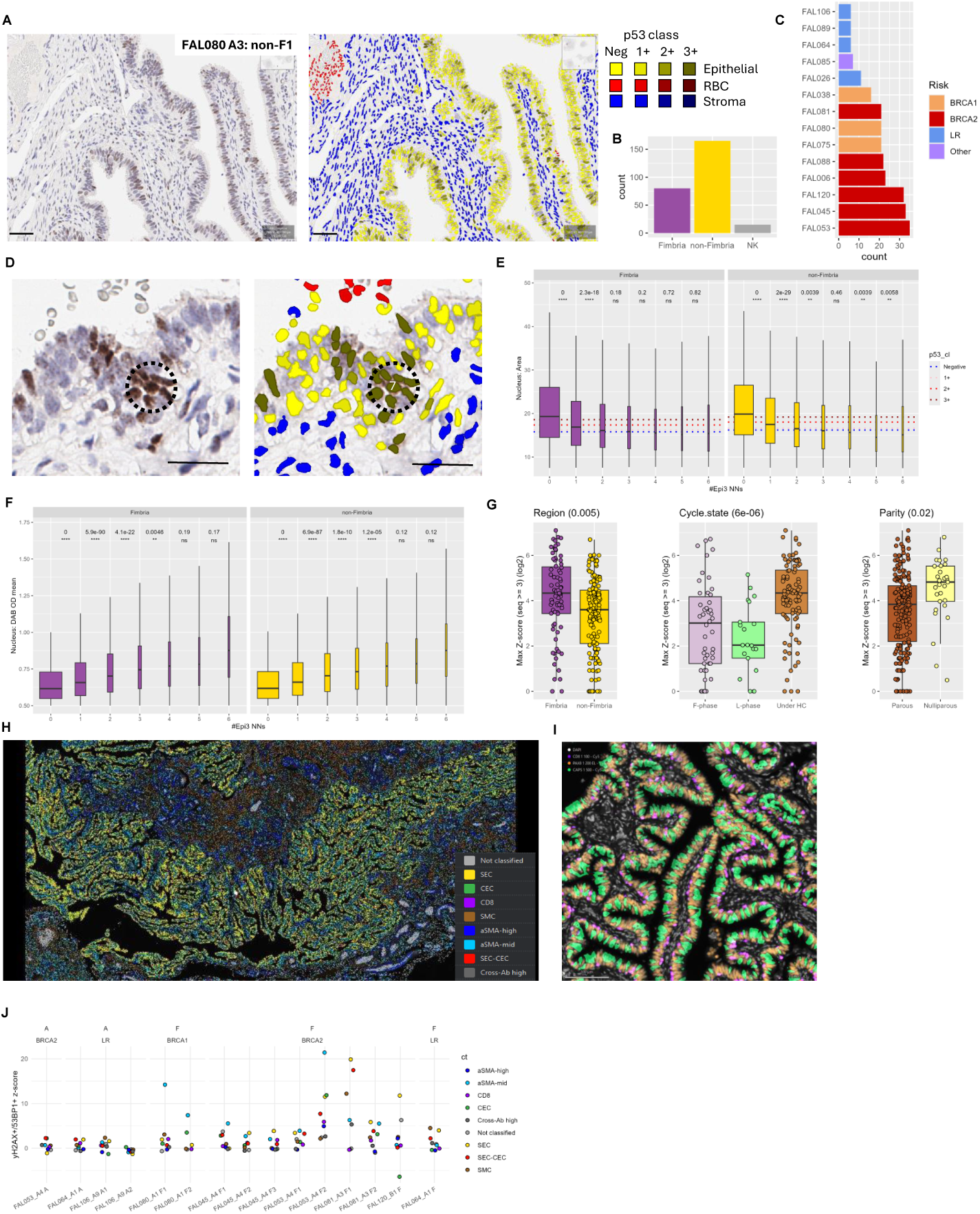
BRCA-associated FT fimbria show cellular changes driven by replication stress. **a,** Representative example of p53 immunohistochemical analysis showing a sub-region within FAL080 (slide A3, non-fimbrial region number 1), using 10x objective lens; scale bar length: 50µm). Legend for right-hand image shows classification for cell-type and p53 expression score. **b,** Bar plot showing total number of regions profiled with p53 immunohistochemistry. NK indicates ambiguous regional location. **c,** Bar plot showing total number of regions analyzed for each subject by risk. **d,** Representative example of an epithelial-3+ streak, with p53 immunohistochemistry staining on the left (using 40x objective lens; scale bar length: 50µm) and cell classification on the right. Dashed circle indicates a cluster of epithelial-3+ cells. The central cell has a streak length of length 7 (annotated in right panel) as the maximal number of epithelial-3+ nearest neighbours is 7. **e,** Boxplots showing nuclear size of epithelial-3+ cells stratified by streak length ≤ 6 and FT region. *P* values indicated are from Mann-Whitney test for adjacent streak lengths. Dotted lines indicate the median nuclei area for all epithelial cells by p53 expression score. **f,** Boxplots showing p53 intensity of epithelial-3+ cells stratified by streak length ≤ 6. **g**, Boxplots showing streak-length Z-score by FT region, cycle-state and subject parity. Panels show *P* values from FDR corrected Kruskal-Wallis test. **h,** Representative example of a multiplex sequential immunofluorescence imaging (mIF) of epithelial fimbrial region from FAL080 A1. Legend indicates cell type classification. **i,** Representative example of FT fimbria showing spatial organization of SECs (PAX8), CECs (CAPS) and CD8 cells (CD8); scale bar length: 100 µm. **j,** Dot plot showing co-occurrence of γ-H2AX and 53BP1 dots by cell-type. Data are ordered by region then subject. Panels indicate risk category.

The overall frequency of p53 positive epithelial cells did not vary by region or risk status (Supplementary Fig. 22d). Larger nuclear size, as a surrogate for S-phase, was associated with higher wild-type p53 expression, with gradual increase in intensity in epithelial - but not stromal - cells (Supplementary Fig. 22e). For each p53-high (3+) epithelial cell, we counted the number of consecutive nearest neighbouring epithelial p53-high cells to spatially detect clusters of cells with high p53 expression (Fig. 5d). In isolated p53-high epithelial cells (e.g. nearest epithelial cell is not p53-high), putative wild-type p53 cells undergoing replication in S-phase, we observed larger nuclei and lower p53 intensity (Fig. 5e, f) compared with longer streaks of p53-high epithelial cells, which are likely to represent patches of p53 mutant cells, an early manifestation of previously defined p53 signatures. The two longest streaks of p53 positive cells identified by our automated methods, were retrospectively classified by our pathologist as STIC and p53 signatures (Supplementary Fig. 23a). We compared the observed cell count on each streak to the expected number (based on the number of p53-positive 3+ epithelial cells and the total number of epithelial cells in the section), yielding a z-score per streak-length and section (Supplementary Fig. 23b). Streak z-score was not associated with *BRCA1/2* status, menopause state or past use of hormonal contraception, but was elevated in the fimbria, in pre-menopausal women using hormonal contraception and in nulliparous women (Fig. 5g and Supplementary Fig. 23c). When looking at the overall z-scores per patient in the fimbria, the putative site of origin for HGSOC, normalised by the non-fimbrial z-scores from the same patient, we observed a fimbrial enrichment for these streaks in *BRCA1/2* subjects and depletion in pre-menopausal women using hormonal contraception, linking these cellular features with known epidemiological HGSOC risk modulators (Supplementary Fig. 23d) and supporting other new evidence suggesting the existence of multiple early p53 signatures expanding at a slow pace in the FT ^51^.

Multiplexed immunofluorescence was performed on six slides using a panel of lineage markers (CD8, PAX8, CAPS, Desmin, α-SMA) and well as markers of cell-cycle arrest and DNA double-strand breaks (p53, γ-H2AX, 53BP1). Cells were annotated as SECs, CECs, SEC-CEC double-positive, CD8, SMCs, α-SMA-high (PV surrounding vessels) and α-SMA-mid cells (scattered fibroblasts) (Fig. 5h–i; Supplementary Fig. 24a and Supplementary Fig. 25). Following RBCs exclusion, based on their high non-specific cross-antibody intensity and localisation (Supplementary Fig. 24b), we defined epithelial neighbourhood in each slide by expanding high-density epithelial cells to their 750 K-nn cells, allowing us to measure cell-type frequency in an unbiased way (Supplementary Fig. 24c). We observed depletion of SECs and an increase in the frequency of α-SMA-high and SMCs upon menopause, and an overall depletion of SMCs in the fimbria (Supplementary Fig. 25d). Detection of wild-type p53 expression is not as sensitive in immunofluorescence staining as it is in immunohistochemistry, as there is no enzymatic signal amplification. We have therefore assessed DNA double-strand breaks as a readout of DNA damage associated with replication stress, by quantifying γ-H2AX and 53BP1 foci in the nuclei through the *COMET* dot detection tool (Supplementary Fig. 26a) ^52,53^. γ-H2AX protein was previously detected in p53-signatures and in normal FTs ^54^. Epithelial cells were enriched with dots of both proteins, with SECs showing stronger γ-H2AX enrichment and CECs a stronger 53BP1 one (Supplementary Fig. 26b). We observed enrichment of double-positive cells for γ-H2AX and 53BP1 mainly in the fimbrial SECs, CECs, and α-SMA-mid cells of *BRCA1/2* subjects (Fig. 5j), with other FT regions not showing the same enrichment. Strikingly, all 3 subjects previously showing the highest histone expression across SECs with elevated *TP53* (Fig. 3h), high number of SNV clones (Supplementary Fig. 19e) and cell-cycle activity within SCNA clones (Fig. 4i) show the highest enrichment for co-detection of γ-H2AX and 53BP1 dots. Overall, these data indicate increased replication stress and associated DNA double-strand breaks in BRCA-associated FT fimbria and support a haplo-insufficient phenotype for BRCA genes in the FT.

## Discussion

Efforts to develop prevention strategies for high-grade serous ovarian carcinoma (HGSOC) have been significantly hindered by the inability to accurately measure efficacy of interventions, owing to the rarity of serous tubal intraepithelial carcinoma (STIC) lesions and the lack of molecular physiology data from high-risk fallopian tubes. Previous bulk transcriptomic and epigenetic analyses of *BRCA*-associated fallopian tubes have reported molecular alterations in high-risk tissues; however, many of these signatures appear confounded by age, surgical indication, and variation in cellular composition ^55^. When correcting for those parameters, no differences were observed between *BRCA*-associated and wild-type FTs ^56,57^. Single-cell analyses have so far been limited mainly to low-risk individuals, constrained by the diagnostic need to preserve tissue integrity and to prevent harm from missed diagnosis of rare STIC lesions.

We integrated over 160,000 single-cell transcriptomes from both *BRCA1/2* mutation carriers and low-risk women to provide a comprehensive single-cell atlas of the human fallopian tube across genetic risk groups, hormonal states, and anatomical regions. This atlas delineates how epithelial cell states, stromal and immune microenvironments, and genomic instability converge to create a landscape of early vulnerability to malignant transformation.

We hypothesized that oestrogens and progesterone could induce proliferative and differentiated cell states in the FT, respectively, in a similar fashion to physiological effects in the endometrium. Under this model, cyclic epithelial shedding during menstruation may serve a protective function by facilitating the removal of chromosomally unstable cells and leading to a lower incidence of serous carcinomas in the endometrium, when compared to the fallopian tubes.

Hormonal exposures further remodel the transcriptional landscape of the fallopian tube in ways that may modulate cancer risk. Menopause depletes ciliated cell populations and induces broad stress-response programs, whereas menstrual cycle stage and use of hormonal contraceptives dynamically reshape both epithelial and stromal compartments (Fig. 2). We identify a luteal-phase–restricted glandular epithelial subset that shares features with luteal-phase endometrial epithelium. Notably, prior use of hormonal contraceptives leaves durable transcriptional imprints across multiple cell types and alters tumour-associated macrophage frequencies, indicating a long-term influence of hormonal modulation on the fallopian tube microenvironment. These findings provide a mechanistic framework that reconciles epidemiological links between ovulation, pregnancy, and hormonal contraception with altered HGSOC risk.

The detection of tumour-associated macrophage–like subsets and cytotoxic lymphocytes within histologically normal fallopian tubes suggests ongoing immune surveillance. Our observed increased expression of MHC-class 2 genes in secretory epithelial cells, with associated activation of immune responses, especially in the context of progesterone-based hormonal contraception, also show hormones may have a role in this immune–epithelial crosstalk. These interactions may determine whether aberrant clones are eliminated or persist, raising the prospect that enhancing early immune clearance could serve as a preventive strategy that preserves fertility.

We observed extreme enrichment of *PLCG2* and *MTRNR2L12* in both *BRCA1* and *BRCA2* carriers, in subjects actively using hormonal contraception and in pre-menopausal ampullar SECs and CECs, whilst there was depletion of the expression of these genes in epithelial cells with glandular phenotype. PLCG2 (1-Phosphatidylinositol-4,5-bisphosphate phosphodiesterase gamma-2) generates 1D-myo-inositol 1,4,5-trisphosphate and diacylglycerol, which influence cellular proliferation, endocytosis, and calcium flux and dysfunction is associated with a variety of diseases including cancer, neurodegeneration, and immune disorders. From its key role as a signalling enzyme in intracellular calcium homeostasis, our data may suggest upregulated crosstalk between cell types in high-risk FT ^29^.

Consistent with prior observations, SECs emerge as the epithelial population most predisposed to malignant progression ^58,59^. These cells preferentially accumulate single-nucleotide and copy-number alterations (Fig. 4) and contain a rare proliferative subpopulation with high *TP53* expression characterized by replication stress, histone–ribosomal imbalance, and transcriptional plasticity (Fig. 3). This “Histone-high/*TP53*-high” state exhibits features of dedifferentiation, G1/S or G2/M checkpoint arrest, and inflammatory cell death accompanied by innate immune recruitment (Fig. 3). These findings support a model in which aberrant clones arising within the fallopian tube epithelium are normally eliminated through *TP53*-dependent surveillance. However, in the context of *TP53* mutation, which is present in histologically normal epithelia ^60,61^, some aberrant cells may evade these checkpoints, allowing persistence of aneuploid clones that accumulate further chromosomal instability. This mechanism is consistent with observations of extensive genomic complexity in early neoplastic lesions ^59^ and with recent evidence of immune evasion following loss of *TP53*-mediated clearance ^54^.

Integration of scRNA-seq–derived instability signatures with DNA-based validation strengthens the view that genomic aberrations in the ostensibly normal fallopian tube are common but usually pruned ^62^, and that persistence of aberrant SECs may be the pivotal event in malignant initiation. The detection of aneuploid clones within histologically normal fallopian tubes suggests that certain chromosomal alterations are tolerated, similar to the viability observed in trisomies 13, 18, and 21. Moreover, the presence of shared chromosomal alterations across both secretory and ciliated epithelial subpopulations indicates that not all genomic aberrations necessarily confer pre-malignant potential but may instead reflect physiologic tolerance to copy-number imbalance.

Our findings of enrichment of aberrant SECs in *BRCA1/2* carriers supports a model in which haploinsufficiency-driven replication stress acts as an early determinant of malignant susceptibility, analogous to mechanisms described in *BRCA*-associated breast epithelia ^44^. Tissue specificity in hereditary cancer syndromes may therefore result from the intersection of epigenetic effects on replication and DNA repair pathways with the intrinsic proliferative dynamics of individual tissues.

Together, these findings position the fallopian tube epithelium of *BRCA1/2* mutation carriers as a tissue under chronic proliferative and replicative stress, in which rare epithelial states foreshadow malignant initiation. By linking genetic predisposition, hormonal context, and cell-intrinsic instability, this work provides a unified framework for developing preventive interventions that target early epithelial vulnerabilities, potentially reducing ovarian cancer risk without surgical intervention.

## Methods

### Ethical approval and clinical data collection

Clinical data and tissue samples for the subjects were collected on the prospective cohort study name of the study (TARGET FAL01: Translational Analysis and Research in Gynaecological Epithelial Tissues - Fallopian Tubes) and was approved by the Institutional Ethics Committee (REC 18/NI/0189). Subjects provided written, informed consent for participation in this study and for the use of their donated tissue for the laboratory studies carried out in this work. Pseudonymised clinical data for all the subjects is provided in Supplementary Information.

### Fallopian tube tissue processing

Fallopian tube tissue was cut into small pieces using a scalpel. Some samples were frozen at this point using CryoStor® CS10 solution. Samples were incubated in DMEM/F-12 containing collagenase and hyaluronidase for 45 min in an orbital shaker. The tissue-mixture was transferred into a 50 ml Falcon tube and centrifuged at 300 g for 5 minutes. Supernatant was removed and the pellet was resuspended in 1ml of Trypsin for 4 minutes; then washed in DMEM/ F-12 and centrifuge at 300 g for 5 min. Supernatant was removed and the pellet was resuspended in 1 ml of dispase for 4 minutes, then washed in DMEM/F-12 and centrifuged at 300 g for 5 minutes. Supernatant was removed and cell pellet was resuspended in 1 ml PBS containing 0.05% albumin (PBSA). Cells were filtered through a 40 µM filter and counted. A 43 µl single cell solution of 372 cells/µl in PBSA were used for single-cell sequencing. When using frozen tissue, the samples were quickly thawed, washed in cold PBS and processed as fresh. Reagents used were;

Collagenase at a final concentration of 1mg/ml (Roche cat no. 11088793001)

Hyaluronidase, final concentration 100U/mL (Sigma cat no. H3506)

Trypsin-EDTA, 0.25% (Stem Cell Technologies cat no. 07901)

Dispase, final concentration 5mg/mL (Stem Cell Technologies cat no. 07913)

### DMEM/F12 (Gibco cat no. 11514436)

CryoStor® CS10 solution (Stem Cell Technologies cat no. 07930)

### Single-cell RNA sequencing

Single-cell RNA-seq libraries were prepared using the Chromium Single Cell 3′ Library & Gel Bead Kit v3.1 and Chromium Chip G Kit according to the Chromium Single Cell 3′ Reagent Kits v3.1 User Guide (CG000315 Rev C; 10x Genomics). Cell suspensions were loaded onto the Chromium Controller with a target recovery of 10,000 cells per channel when processing individual subject samples. For multi-regional subject samples, multiple specimens were multiplexed prior to 10x Genomics processing, with an expected recovery of up to 18,000 cells per reaction.

Library quality was assessed using an Agilent 4200 TapeStation with High Sensitivity D1000 ScreenTape to evaluate library size distributions and a Qubit 4.0 Fluorometer (Thermo Fisher Scientific; Qubit dsDNA HS Assay Kit) to determine dsDNA concentration. Libraries were normalized and pooled at equimolar concentrations. Pool concentrations were verified by qPCR using the KAPA Library Quantification Kit on a QuantStudio 6 Flex instrument prior to sequencing.

Sequencing was performed on an Illumina NovaSeq 6000 platform using the following read configuration: 28 bp for Read 1, 10 bp for the i7 index read, 10 bp for the i5 index read, and 90 bp for Read 2. Reagents used were;

Chromium Next GEM Single Cell 3ʹ Kit v3.1, 16 rxns (10x Genomics PN-1000268)

Chromium Next GEM Chip G Single Cell Kit, 48 rxns (10x Genomics PN-1000120)

Dual Index Kit TT Set A, 96 rxns (10x Genomics PN-1000215)

SPRIselect Reagent Kit (Beckman Coulter B23318)

DNA High Sensitivity D1000 and D5000 (Agilent, Cat# 5067-5584/5067-5592)

Qubit™ dsDNA HS Assay Kit (Thermo Fisher, Cat# Q32854)

KAPA Library Quantification Kit for Illumina Platforms (KAPA Biosystems KK4824)

Cellranger v7.01 (10x Genomics) was used to generate raw count matrices with the hg reference (refdata-gex-GRCh38-2020-A). Barcodes potentially swapped during sequencing were detected and removed using DropletUtils::swappedDrops ^63^ with min_frac = 0.9. Cells with at least 1500 UMIs from non-mitochondrial genes and less than 20% UMIs from mitochondrial genes were kept for further analysis. Cells from multiplexed runs were assigned to their source sample (see Sample multiplexing below).

### Sample multiplexing

Several samples from multi-regional subjects were multiplexed before 10x processing. De-novo genotyping was used to demultiplex these samples. In each multiplexed pool we had at most one sample from a new subject, the rest were from subjects that other regions of their FT were already sequenced. Demultiplexing followed the processing steps of the non-multiplexed samples until having a set of QC filtered cells. The original 10x BAM files of the sample and previously processed samples from the same subjects were filtered to contain only QC filtered cells and were concatenated into a single BAM file. The merged BAM file was used for de-novo genotyping and clustering by genotyping, both done with *Souporcell* ^64^ using the total number of subjects in the pool as the expected number of clusters. Clusters were assigned to subject based on the cluster membership of cells from the previously processed (non-multiplexed) samples. Inter-subject doublet cells from the multiplexed runs were detected and removed. We had a total of 10 multiplexed runs with 2–4 samples in each run. A single run (run 10) failed and discarded, where the expected clusters were not detected and quality metrics were extremely poor. The rest were clustered perfectly by subject, judging by the perfect cluster assignment of the non-multiplexed samples in the pool (Supplementary Fig. 1).

### Metacell modeling

Extremely strong genes and those prone to technical effects were removed from the count matrix (mitochondrial, immunoglobulins and a few strong non-coding genes, Supplementary Table 4). The cells were then partitioned into metacells in an initial model used for basic quality control and to define the gene sets to be blacklisted from affecting the final model (gene modules correlated with cell cycle, hypoxia, interferon and stress responses, Supplementary Table 4). Scrublet ^65^ was used with 0.35 as a cut-off for doublet score to remove putative doublet cells. The singlet cells were used to create a second metacell model. Broad cell-type signatures (Supplementary Table 5) were computed on these metacells to further detect and remove 8 putative doublet metacells (Supplementary Fig. 1h, i). The final metacell model contained 168,679 cells partitioned into 945 metacells. Cell-type/state was assigned to the filtered metacells by manual annotation of metacell clusters in the hierarchically clustered metacell confusion matrix.

### Intra-cell-type compositional controlled differential gene expression

We aimed to find differentially expressed genes within a cell-type that are truly driven by the selected clinical feature. We performed a pseudo-bulk differential gene expression analysis by pooling together UMIs in each of the two compared groups, while performing these filters and normalisations to ensure the results are robust and not driven by compositional changes or an extreme expression in a single subject:

- Requiring at least 10 cells in at least 3 subjects in each comparison group
- Requiring at least 100 total cells in each comparison group
- Leave-one-subject-out to eliminate subject specific results – remove one subject at a time, repeat the analysis, and only keep the genes differentially expressed in the same direction (enriched or depleted) in all iterations.
- Balancing clinical composition (e.g. by risk, menopause, region): Partitioning the compared samples by the balancing features values, keeping partitions with at least 10 cells in either group, normalise the total number of UMIs within each partition before summing up the normalised total UMI counts to create the pseudo-bulk count.

Supplementary Table 8 summarises the differential analysis runs, the features used for comparison, the features used to restrict the runs (e.g. only in pre-menopausal subjects), and the features used for compositional balancing.

### SNV and SCNA calling

We used *Scomatic* ^47^ to detect SNVs. *Scomatic* is using the supplied cell-type annotation to filter cross-cell-type mutations, assuming that true somatic mutations will be lineage specific. It also filters mutations that appear in an extensive panel of normal the authors created, and suspected events of RNA editing. We ran *Scomatic* per subject, pooling together all the subject’s samples into a single bam file. We supplied high-level cell-type annotation to the tool (Endothelial, Epithelial, Fibroblast-PVC, Immune, Schwann, Erythrocytes), reasoning that detailed annotation will be too restrictive by filtering intra-cell-type mutations. We filtered out suspected germline mutations (VAF = 1) and suspected common SNPs (MAF ≥ 0.0001). We found 11 mutations shared by several subjects which do not seem to be a sequencing technical artefact (they are shared across different 10X and sequencing runs) and filtered those too as suspected SNPs.

We used *Numbat* ^49^ to detect SCNAs. This tool uses the mean expression in genomic bins as a copy-number proxy, but also leverages SNPs haplotype information to improve its copy-number estimation. We ran *Numbat* by subject and cell-type, filtering cell-types with at least 50 cells in a subject. To minimise SCNAs driven by cell-type specific expression, we pooled together cells from the same cell-type across all subjects to be used as the normal reference. We used *Numbat* default parameters, which are tailored for tumour level copy-number detection, and are therefore stringent for our normal tissue dataset.

### Statistics

All statistical analyses were performed in R. Two-group comparisons used the two-sided Mann-Whitney-Wilcoxon rank-sum test (wilcox.test, stats). Comparisons across three or more groups used the Kruskal-Wallis rank-sum test (kruskal.test, stats), which is non-directional. For multiple hypothesis corrections, p-values were corrected by the Benjamini-Hochberg false-discovery-rate procedure (adjust_pvalue, rstatix; p.adjust, stats).

### Single-cell DNA-seq generation and data processing

DLP+ single-cell DNA-seq profiles and downstream data analyses were performed as previously described ^50^, resulting in copy number estimation in 500 kb bins and multiple quality metrics per cell. The following steps were performed to filter good quality cells:

- Quality score ≥ 0.75
- S-phase probability < 0.5
- Supported by ≥ 250,000 mapped reads
- Species identified as human (grch37)
- Outlier cells removed (with mean copy-number further than 2 s.d. from the population median)

The following steps filtered good quality genomic bins:

- Bin mappability ≥ 99%
- Fraction of cells in bins marked as ‘ideal’ by the DLP+ pipeline ≥ 80%

### H&E and immunohistochemistry (IHC)

Haematoxylin and Eosin (H&E) slides were stained according to the Harris H&E staining protocol and using an Epredia™ Gemini™ AS Automated Slide Stainer. Paraffin embedded sections of 3 μm were stained using Leica Bond-III fully automated IHC system. Briefly, slides were retrieved using Bond Epitope retrieval solution 2 for 20 minutes and p53 antibody was applied for 30 minutes. Bond™ Polymer Refine Detection System (Leica Microsystems) was used to visualise the brown precipitate from the chromogenic substrate, 3,3′-Diaminobenzidine tetrahydrochloride (DAB). Slides were scanned at 40× using Aperio AT2 digital pathology scanner system. Antibodies used were; P53 (Leica, Clone DO-7, catalog no. PA0057).

We used *QuPath* ^66^ (v0.5.1) to analyse the p53 IHC slides. We followed the recommended pipeline for processing each slide (full script is available in the paper code repository). Briefly, image type was set to brightfield H-DAB, positive cell detection using ‘Optical density sum’ with p53 cutoffs of 0.15, 0.3 and 0.5 for 1+, 2+ and 3+ p53 positive cells, respectively. A classifier trained on manually annotated epithelial, stromal and RBC/debris cells using examples from all slides was used for cell-type assignment. Each tissue fragment with epithelial cells in every slide was manually annotated by a pathologist as coming from the fimbria, non-fimbria or from and unknown source (NK), and downstream analysis was usually performed on the level of these annotated regions.

### Multiplexed sequential immunofluorescence imaging

Multiplexed sequential immunofluorescence (seqIF) staining and imaging was performed using 3 µm formalin-fixed paraffin-embedded (FFPE) sections that were preprocessed using the PT module with dedicated reagents at pH 9 (Epredia). Subsequently, an automated COMET™ platform (Lunaphore Technologies) was used for the seqIF comprising 5 antibodies (Supplementary Table 12). Primary and secondary antibodies were prepared offline and loaded on the instrument together with proprietary buffers. The X-plex protocol was generated using the COMET™ Control Software and loaded accordingly. The seqIF workflow was performed on 4 slides at a time and consisted of 13 cycles, one cycle to prevent autofluorescence followed by 12 cycles of staining. Each cycle included the following steps: wash, elution buffer 2 min incubation, quenching 30 s incubation, primary antibody 8 min incubation, secondary antibody with DAPI 2 min incubation, wash with imaging buffer and image capture.

Slide images from COMET™ were analysed with the HORIZON™ software (v2.3.0, Lunaphore). Default HORIZON™ parameters were used to detect nuclei based on DAPI (1–99.8% quantiles, extend cytoplasm = 5 px, and measure mean antibody intensity in nuclei, cytoplasm and both). Cell-types were hierarchically assigned to cells based on antibody staining and localization (Supplementary Table 13).

Slide specific intensity thresholds were used (Supplementary Table 14). Slide FAL106_A9 had a non-specific CAPS staining and for that slide CEC classification was moved down after SMCs to avoid misclassification of non-CECs as CECs. To remove RBCs mis-classified to other cell types due to sporadic high antibody intensities, we detected and removed suspected RBCs from downstream analysis. An initial set of putative RBCs was defined by cells in high density that the majority of their nearest neighbours are ‘not classified’. Some slides (FAL064, FAL045, FAL081) had RBCs with multiple extremely high antibodies. We marked cells with at least 6 high intensity antibodies (≥ 95% quantile) as multi-high cells and added them to the initial set of RBCs. We then expanded the initial set of RBCs by 200 iterations of majority k-NN voting (K=25) to mark cells around RBCs as RBCs.

## Acknowledgements

We thank all subjects who participated and donated tissue samples to this study and acknowledge the support of the University of Cambridge, Cambridge University Hospitals NHS Foundation Trust and Cancer Research UK. We thank the Addenbrooke’s Hospital Gynaecological Oncology clinical team and the Breast and Gynaecological Cancer Trials team for recruitment and clinical sample collection. We acknowledge the technical support provided by the Bioinformatics Core Facility (RRID:SCR_028664), Compliance and Biobanking, Genomics Core Facility (RRID:SCR_028699), Histopathology and In Situ Hybridisation Core Facility (RRID:SCR_028183), Microscopy Core Facility (RRID:SCR_028019), Research Instrumentation and Cell Services Core Facility (RRID:SCR_028062) and Scientific Computing at the Cancer Research UK Cambridge Institute. We acknowledge the support provided by the Addenbrooke’s Human Research Tissue Bank (supported by the NIHR Cambridge Biomedical Research Centre NIHR203312). We thank C. Caldas and the Addenbrooke’s Hospital Gynaecological Oncology team for valuable discussions and support of this study.

## Authors contributions

Y.E-L. methodology, software, validation, formal analysis, investigation, data curation, writing - original draft, writing - review & editing.

M.V. methodology, validation, formal analysis, investigation, data curation, writing - original draft, writing - review & editing.

K.K. methodology, investigation, writing - review & editing.

D.K. investigation, data curation, writing - review & editing.

J.H. investigation.

R.C. writing - review & editing, resources.

R.N. investigation, resources.

S.W. methodology, formal analysis, investigation.

S.K. investigation.

S.A. methodology, investigation, resources.

C.S. conceptualization, methodology, investigation, resources, writing - review & editing, supervision.

M.J.L. methodology, investigation.

J.D.B. conceptualization, methodology, investigation, resources, data curation, writing-original draft, writing - review and editing, supervision, project administration, funding acquisition.

F.C.M. conceptualization, methodology, investigation, data curation, writing - original draft, writing - review and editing, supervision, project administration, funding acquisition.

## Data and Software Availability

The accession number for the raw sequence reported in this paper will be EGA: EGAXXX. The processed data can be viewed and analysed interactively at https://ft-scrnaseq.shinyapps.io/ft_scrnaseq_mcview_bundle. Scripts reproducing the analysis are available at https://github.com/Brenton-Group/Fallopian_Tube_scRNAseq

---

**Supplementary Figure 1:**
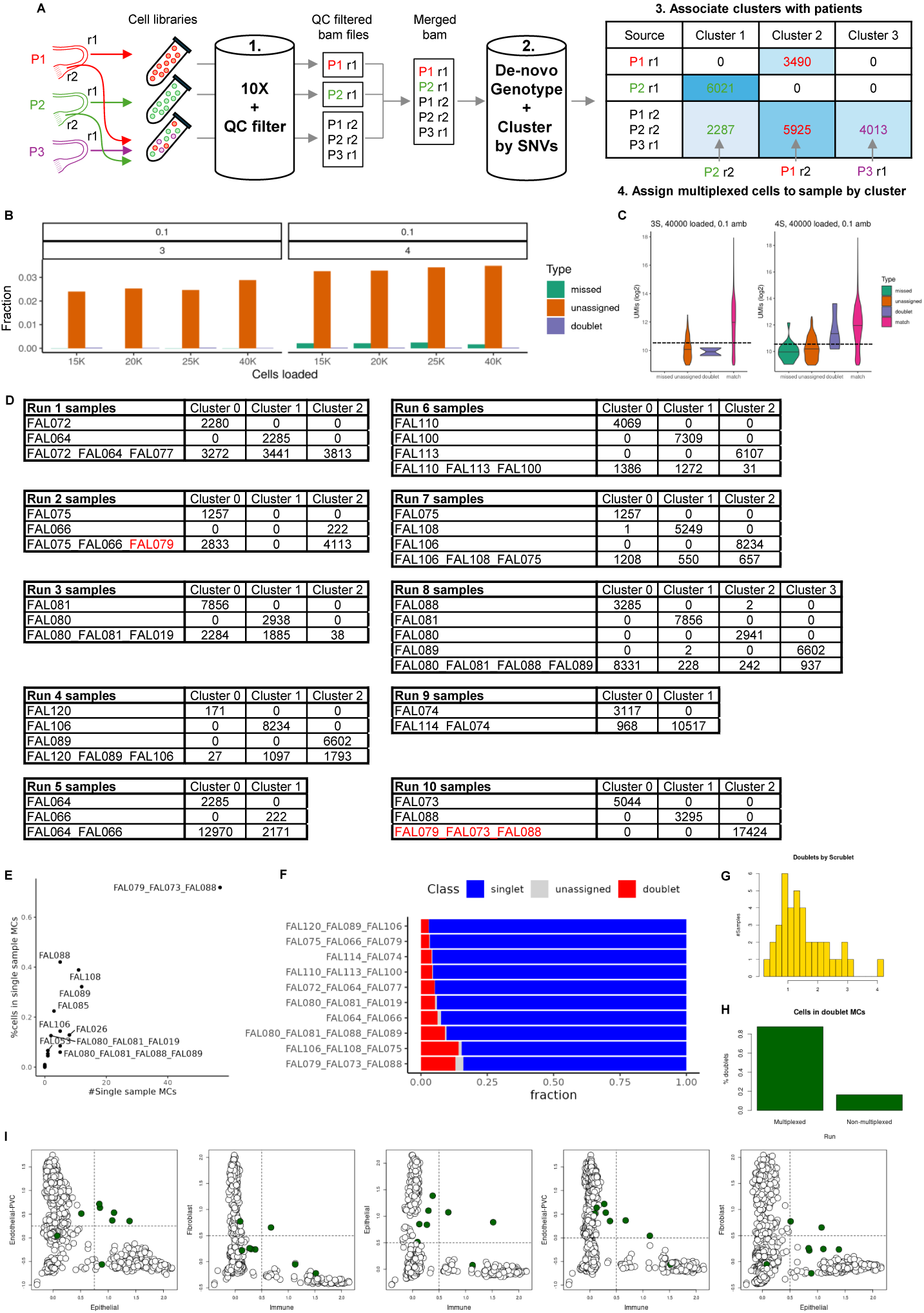
Pipeline and quality control (QC) for sample multiplexing. **a,** Samples multiplexing diagram. The cells of a sample from a region (r1) of one subject (P3) are pooled before 10x processing with cells from other subjects (P1 r2, P2 r2) that have samples from other regions (P1 r1, P2 r1) that were previously processed. After 10x processing, the reads of cells passed quality control from all 3 runs are marked for their origin run and pooled into a merged bam file. This file is processed by *Souporcell* ^64^ for de-novo genotyping and clustering by genotype, with the parameter specifying the expected number of clusters matching the number of subjects in all samples. The clusters are associated with subjects by the membership of the cells from the non-multiplexed runs (P1 r1, P2 r2) which allows the assignment of cells in the multiplexed run to their sample of origin. **b**, Simulation of multiplexed runs to test de-multiplexing performance. Reads from either 3 or 4 non-multiplexed samples were pooled to simulate a multiplexed run with varying number of cells loaded to the 10x platform (15,000 to 40,000). Fraction of doublets was estimated by the number of loaded cells and were simulated by switching one cell barcodes to another one. Ambient noise was simulated by randomly barcode switch on the read level with low (5%) and high (10%) ambient noise level (results for 10% ambient noise are shown). See Supplementary Table 6 for full simulation results. Bar plots show the fraction of unassigned, wrongly-assigned-as-doublets or mis-classified cells. Only misclassification to another sample generate a de-multiplexing error, as cells that are unassigned or wrongly assigned as doublets are safely discarded. Misclassification only appears when multiplexing 4 samples (showing here data for 10% ambient noise) and with extremely low frequency. **c,** Violin plots showing number of UMIs/cell by cell classification in the two most extreme subjects (maximal cell loading and high ambient noise). Dashed line marks the minimal number of UMIs we used to filter good quality cells (1500 UMIs) and shows that no misclassification of good quality cells occurs when multiplexing 3 samples. **d,** Cells cluster membership tables of all 10 multiplexed runs. Non-multiplexed runs that are used to associate a cluster with a subject are on the top rows. Perfect classification can be estimated by the perfect assignment of the non-multiplexed samples. No cells were assigned to FAL079 in run 2, probably due to lack of viable cells from that sample. Accordingly, the run that multiplexed another region from FAL079 (run 10) failed too with no cells clustering with the non-multiplexed samples. **e,** A single-sample metacell was defined as having more than 95% of its cells from a single. The failed multiplexed run 10 (FAL079_FAL073_FAL088) had an extremely high number of single-sample MCs that contained most of its cells (y-axis), suggesting the sample suffered from a severe technical effect. **f,** Classification results for the multiplexed runs, only singlet assigned cells are maintained. **g,** Percentage of cells assigned as doublets by *Scrublet* ^65^ per sample. **h,** Percentage of cells from cross-cell-type manually detected doublet MCs in multiplexed and non-multiplexed runs. Doublet frequency in multiplexed runs is expected to be higher due to the higher number of loaded cells in these runs. **i,** Broad cell-type signature scores across all MCs. Dashed lines mark cutoffs used to define the doublet MCs (coloured in green).

**Supplementary Figure 2:**
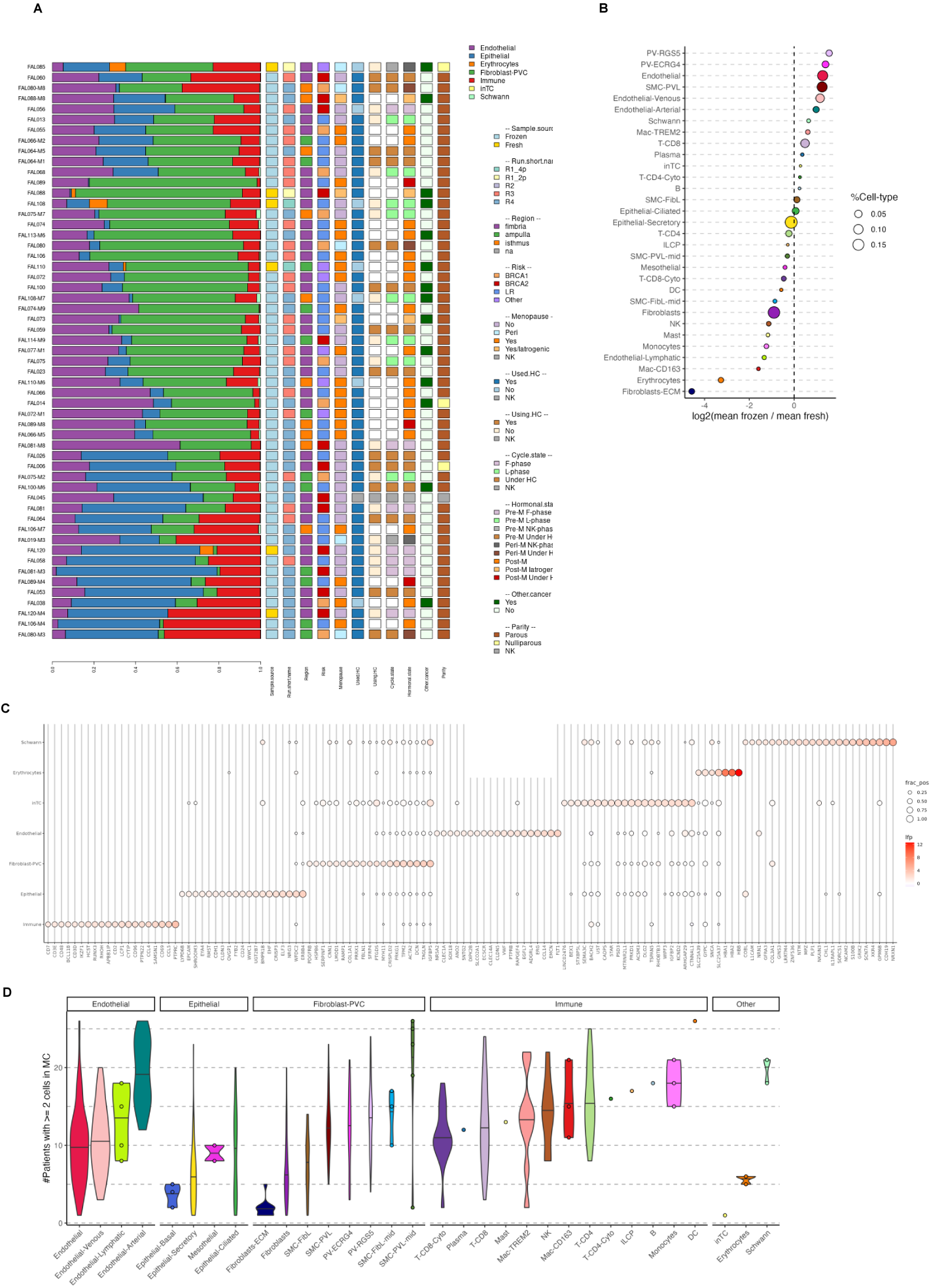
Sample high level composition and metacell annotation. **a,** Samples high-level cell-type composition. Clinical features are shown on the right, samples are clustered by their composition profile. All samples have cells from the main cell-types, although in varying frequencies related to sampling. **b,** Plot depicting comparison between the mean prevalence of each cellular population in fresh and frozen samples **c,** High-level cell-type gene markers heatmap with fraction of positive cells as circle size and gene enrichment as its colour. **d,** Boxplot showing the number of subjects contributing cells to a metacell (≥ 2 cells) by detailed cell-type, black line is the median and whiskers are IQR times 1.5, grouped by high-level cell-types.

**Supplementary Figure 3:**
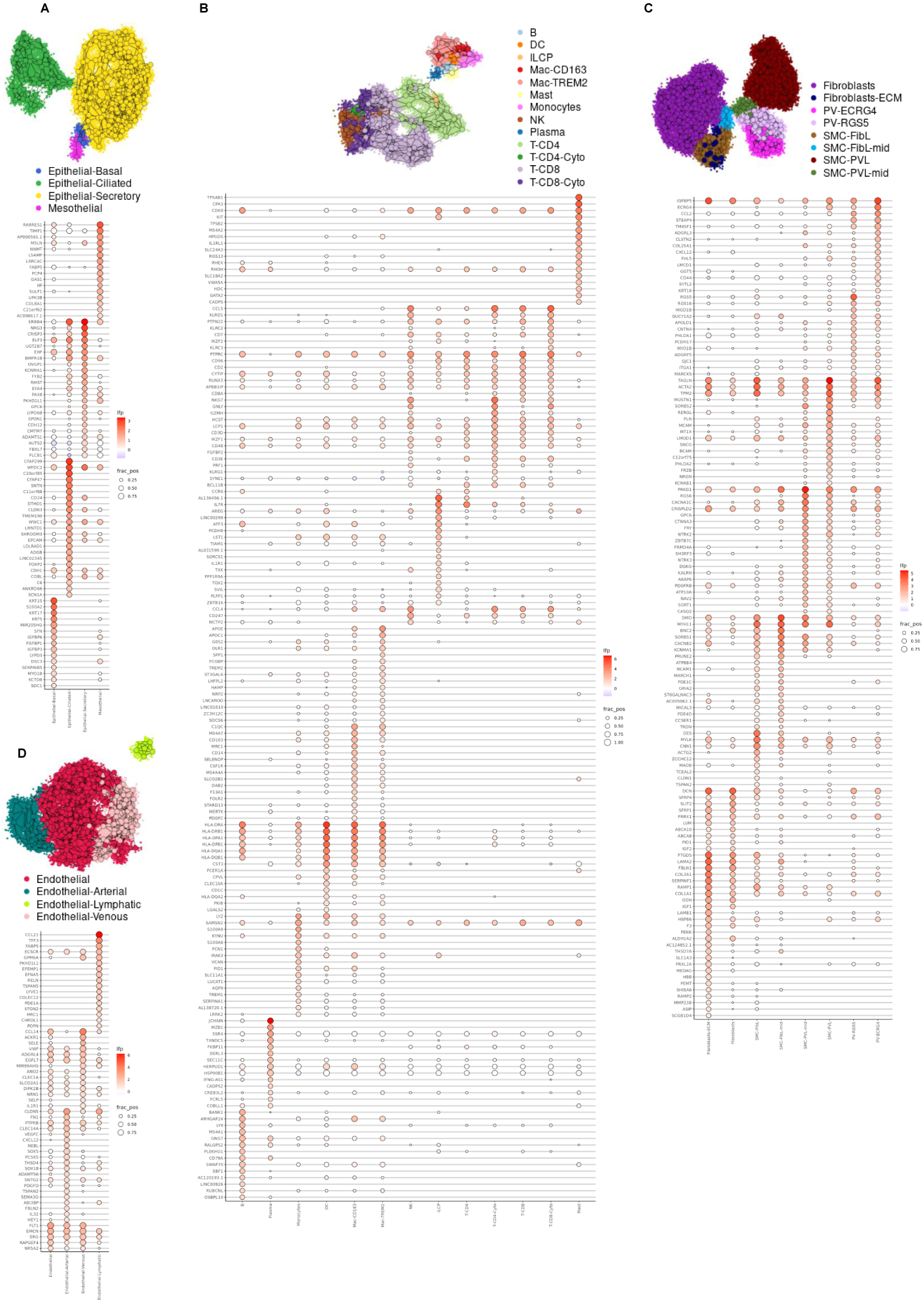
Detailed metacell annotation. **a,** Cell-type annotated 2D map of the 197 epithelial metacells and key cell-type specific genes heatmap with fraction of positive cells as circle size and gene enrichment as colour. **b,** Cell-type annotated 2D map of the 127 immune metacells. **c,** Cell-type annotated 2D map of the 416 fibroblasts/PVC metacells. **d,** Cell-type annotated 2D map of the 201 endothelial metacells.

**Supplementary Figure 4:**
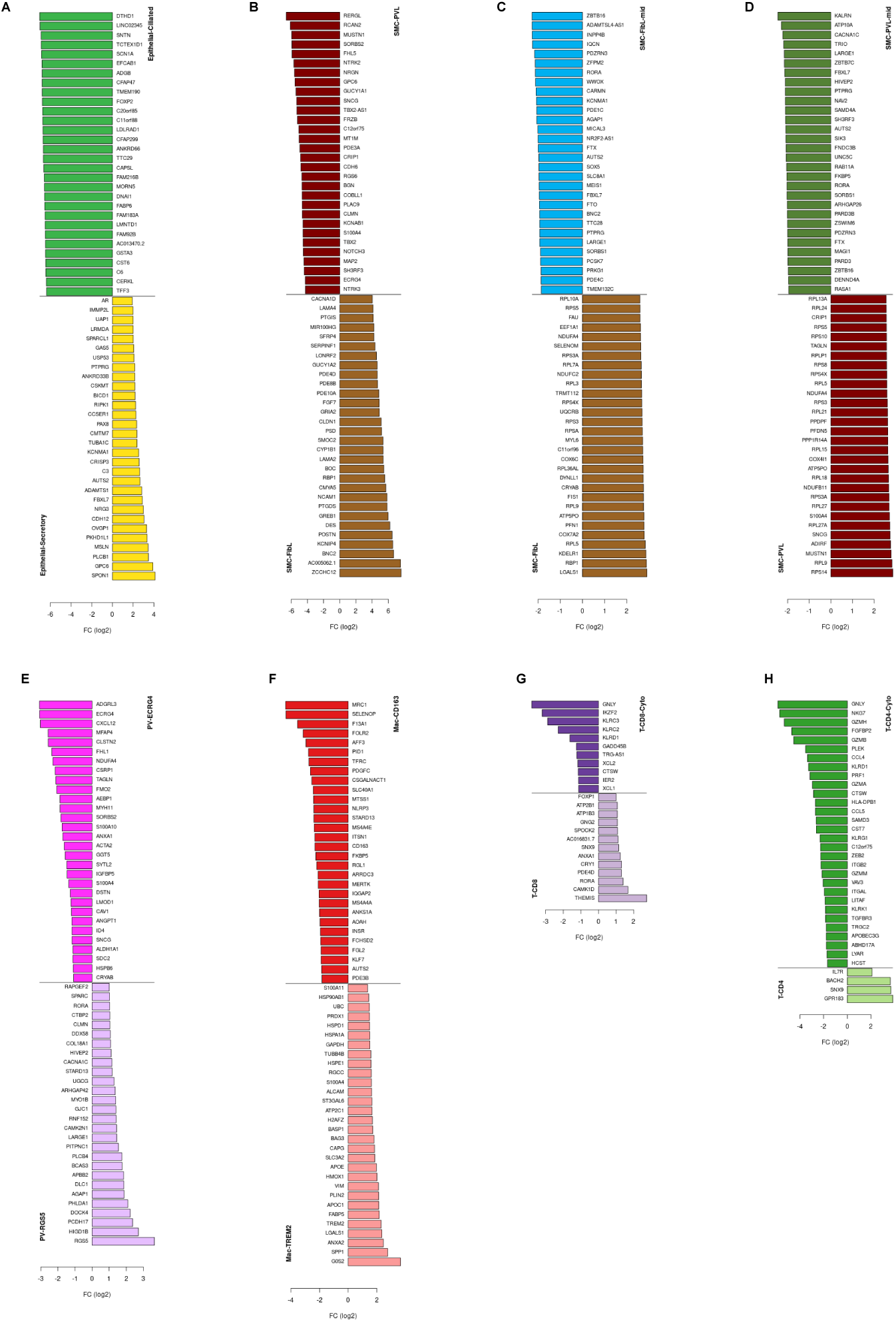
Top differentially expressed genes for pairwise comparison of cell-types. **a,** Epithelial-Secretory vs Epithelial-Ciliated. **b,** SMC-FibL vs SMC-PVL. **c,** SMC-FibL vs SMC-FibL-mid. **d,** SMC-PVL vs SMC-PVL-mid. **e,** PV-RGS5 vs PV-ECRG4. **f,** Mac-TREM2 vs Mac-CD163. **g,** T-CD8 vs T-CD8-Cyto. **h,** T-CD4 vs T-CD4-Cyto. Genes were filtered with LFC ≥ 1 and *P* ≤ 1e-20.

**Supplementary Figure 5:**
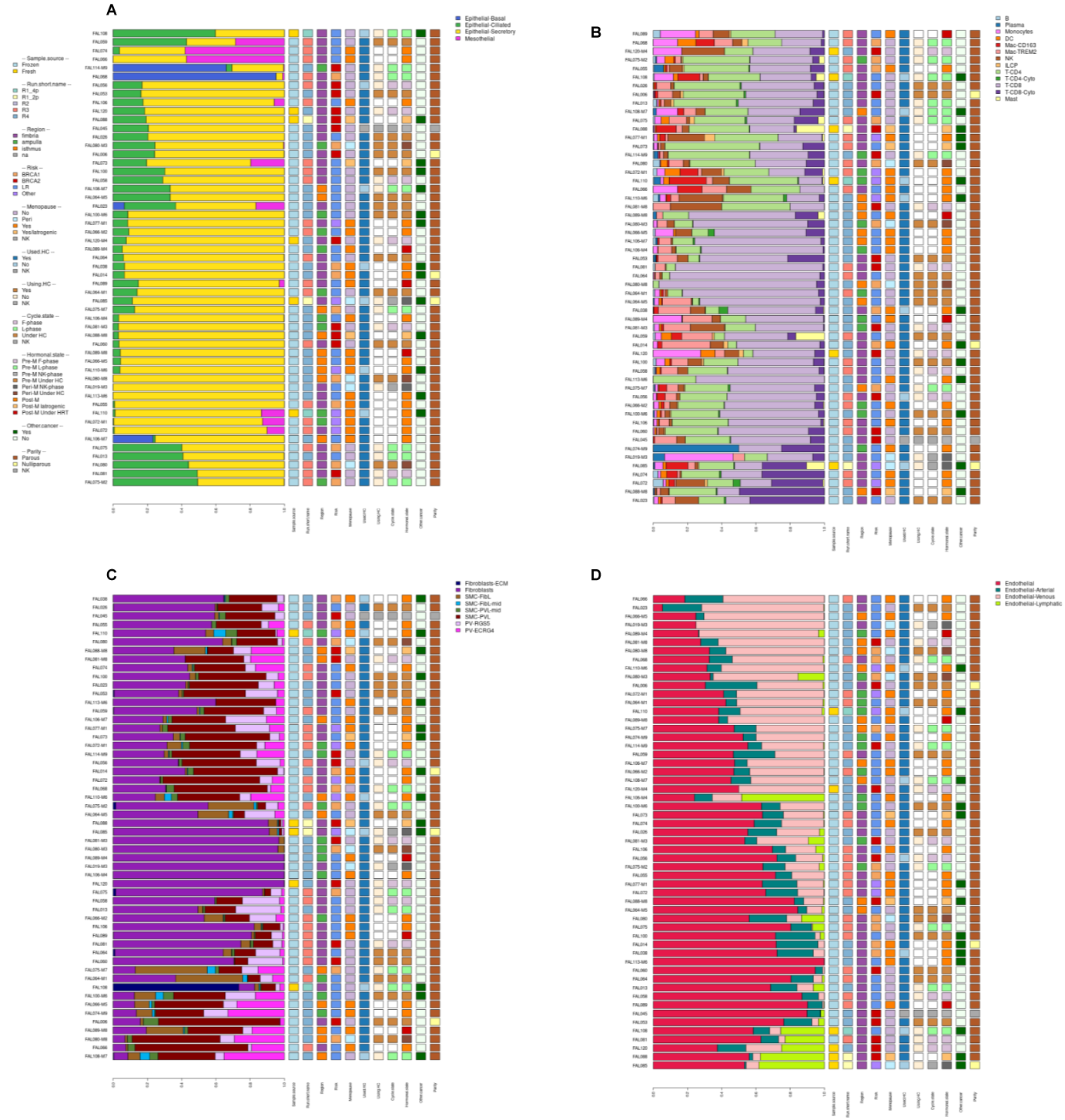
Cell type detailed composition of samples. **a,** epithelial cell type. **b,** immune cell type. **c,** fibroblasts/PVCs cell type. **d,** endothelial cell type.

**Supplementary Figure 6:**
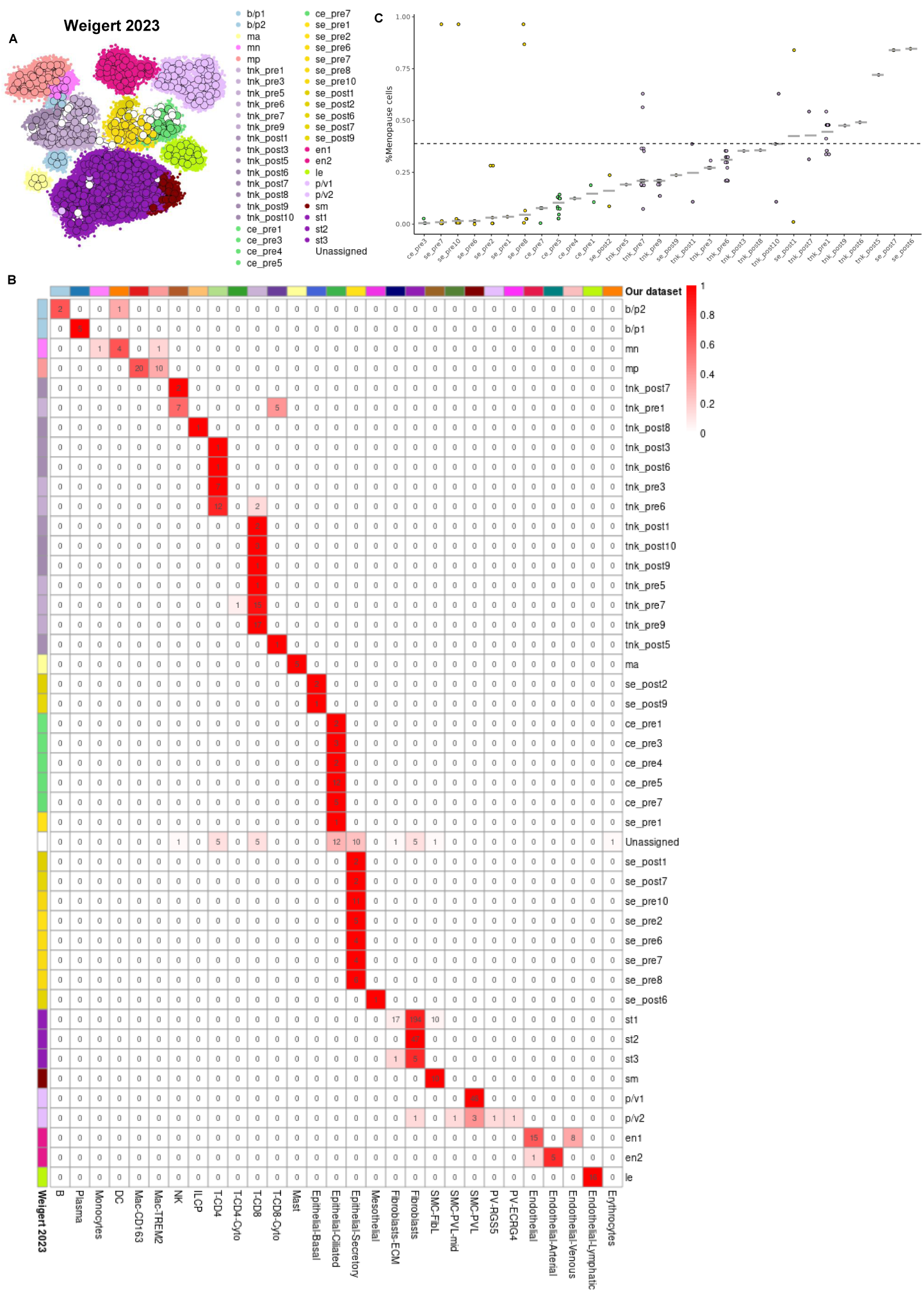
Metacell analysis of published dataset with 10 pre-menopausal and 7 post-menopausal FTs. **a,** Cell-type annotated 2D map of the 614 Weigert et. al^2^ metacells with the cell-type colour legend. Processing of the published scRNA-seq count matrix with *metacell* partitioned 119,279 cells into 614 metacells. We applied a majority vote among the cells in each metacell on the authors original annotation to annotate the metacells. A total of 41 metacells without a dominant cell-type (majority vote < 50% of cells) were classified as unassigned. **b,** Comparison between Weigert at. al metacells (rows) and the metacells in this manuscript. Each of the published metacells is paired with the top correlated metacell from this dataset, showing the clustered pairs count matrix. **c,** Percentage of cells from post-menopause subjects in each metacell, grouped by authors derived cell-type, demonstrating the granularity of the metacells.

**Supplementary Figure 7:**
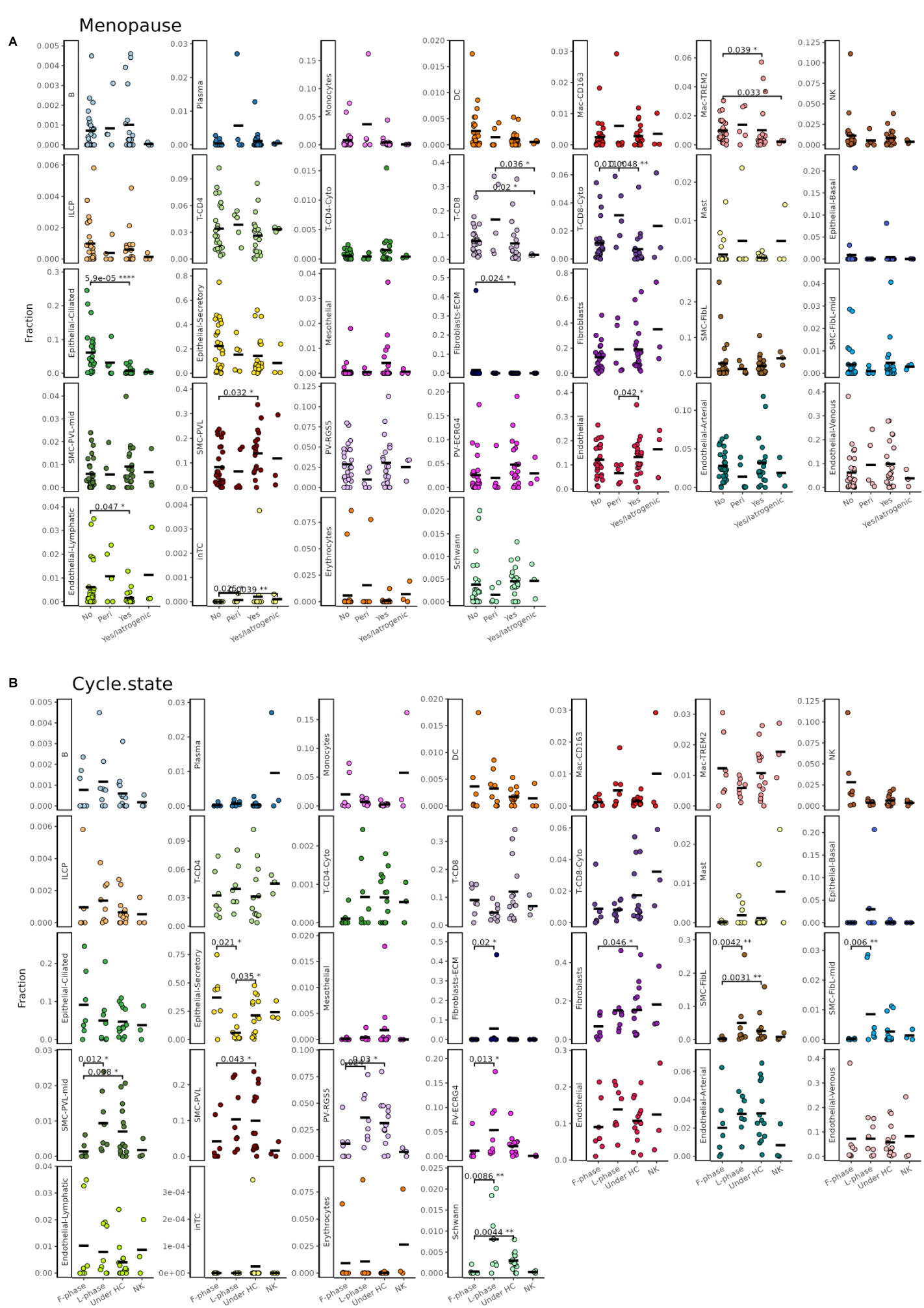
Cell-type abundance comparison across samples. **a,** Menopause state **b,** Menstrual cycle state. P-values by pairwise Mann-Whitney test within each cell-type are shown. Only CECs in menopause are statistically significant by FDR corrected Kruskal-Wallis test across all cell-types.

**Supplementary Figure 8:**
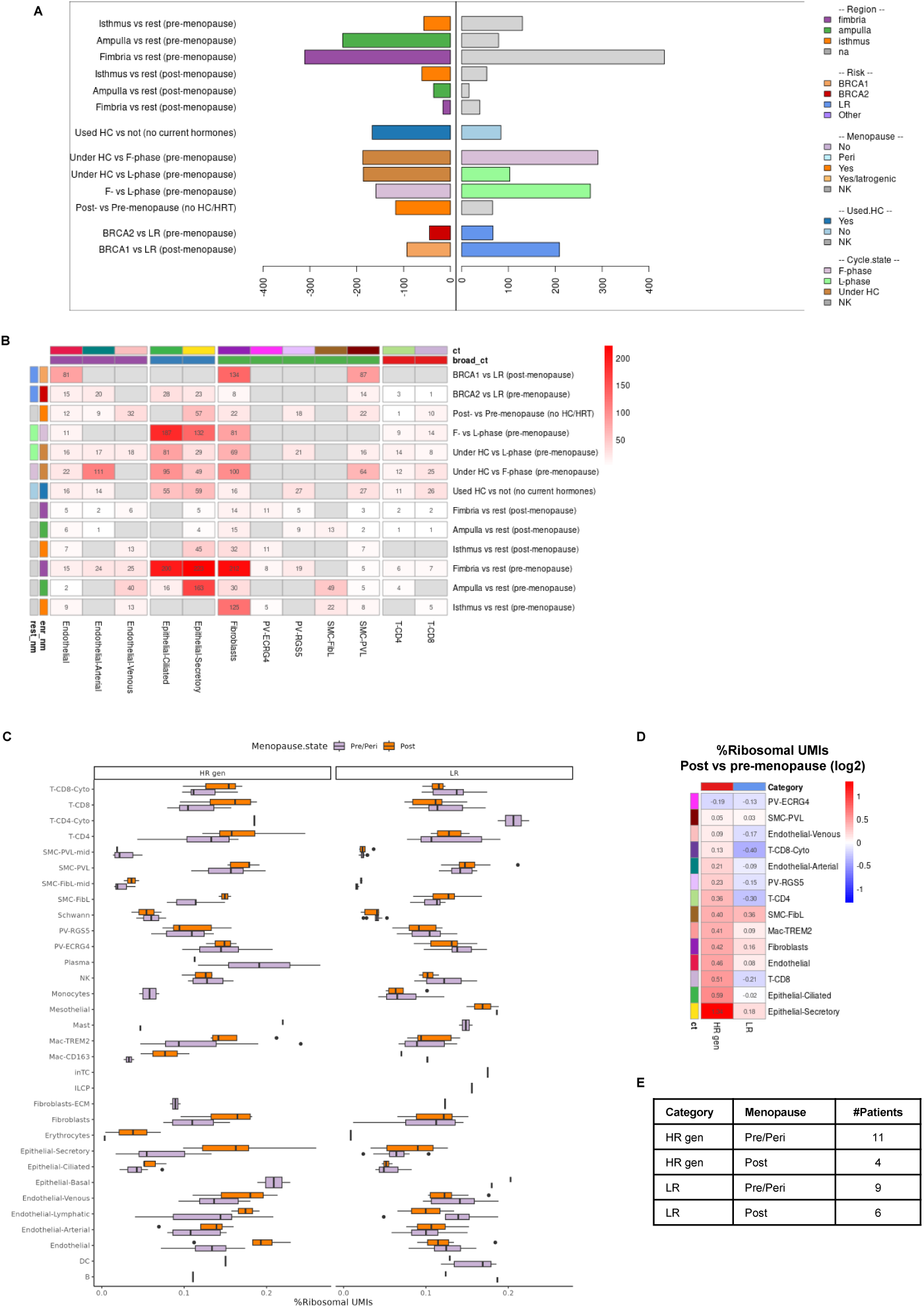
Differential expressed genes (DEG) across clinical categories. **a,** Total number of DEGs across all cell-types per intra-cell-type DE comparison, enriched genes are shown to the left, depleted on the right **b,** Total number of DEGs (enriched and depleted) by cell-type and DE comparison. **c,** Fraction of UMIs from ribosomal genes in a sample per cell-type, comparing pre/peri-menopausal to post-menopausal subjects, separating LR subjects from *BRCA1* and *BRCA2* (HR gen) subjects. **d,** Enrichment of ribosomal gene UMIs upon menopause, showing the log2 ratio of post-vs pre/peri-menopause values per cell-type and risk category. Cell-types are ordered by the ratio in high-risk subjects. **e,** Subject count by broad risk category and menopause status.

**Supplementary Figure 9:**
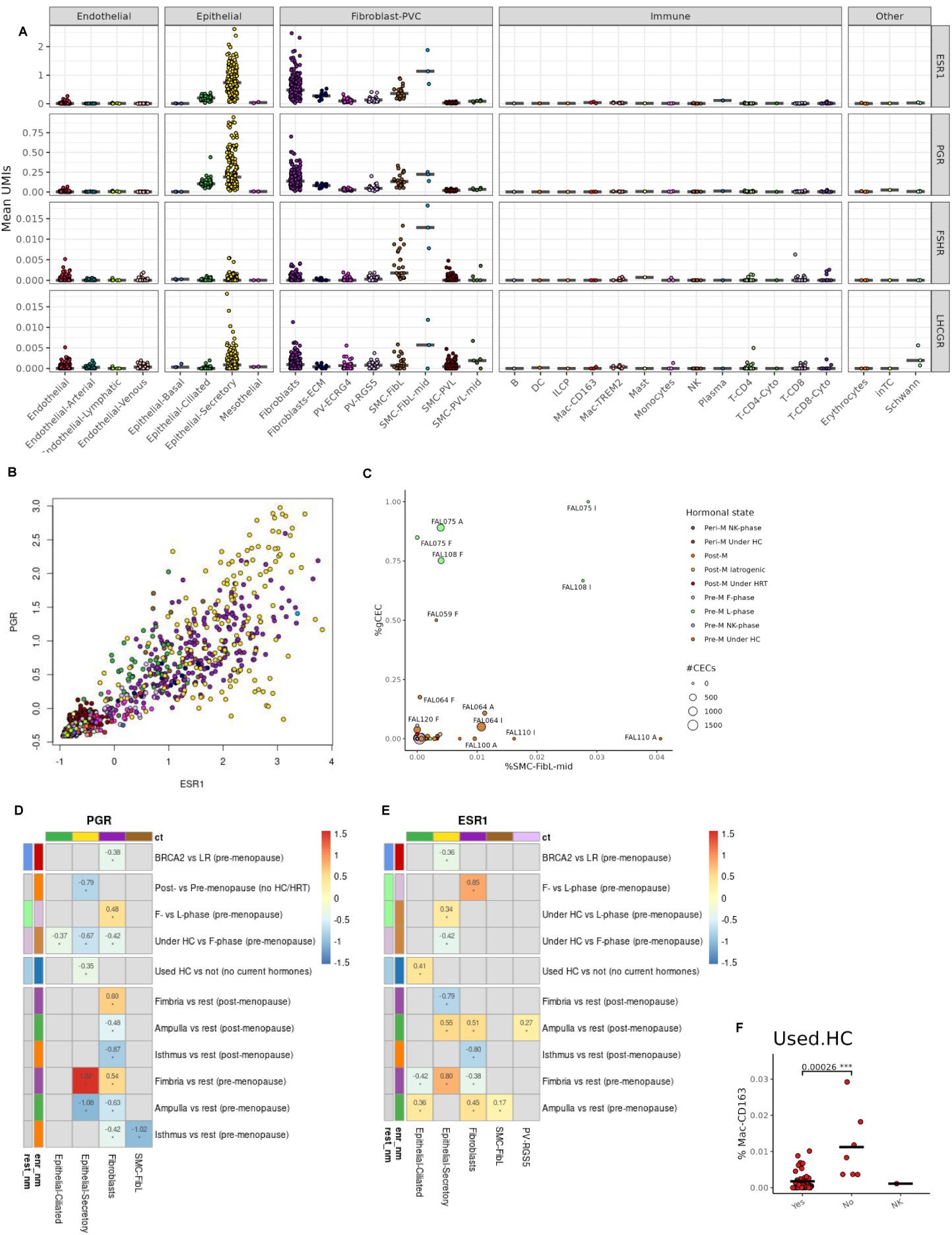
Expression of hormonal receptors across metacells and clinical categories. **a,** Expression (count per 1000 UMIs) of hormone receptor genes across all metacells. **b,** Gene enrichment of PGR and ESR1 in metacells show their coordinated expression. **c,** Fraction of gCECs vs fraction of SMC-FibL-mid cells per sample, coloured by hormonal state (combining menopause and hormone usage), circle size mark number of cells, subject and FT region are shown for high value samples. **d,** All cell-type and DE analysis runs where PGR had an absolute LFC ≥ 1. **e,** All cell-type and DE analysis runs where ESR1 had an absolute LFC ≥ 1. **f,** Fraction of Mac-CD163 cells in samples by their past HC usage status (never used vs used), *P* value by Mann-Whitney test.

**Supplementary Figure 10:**
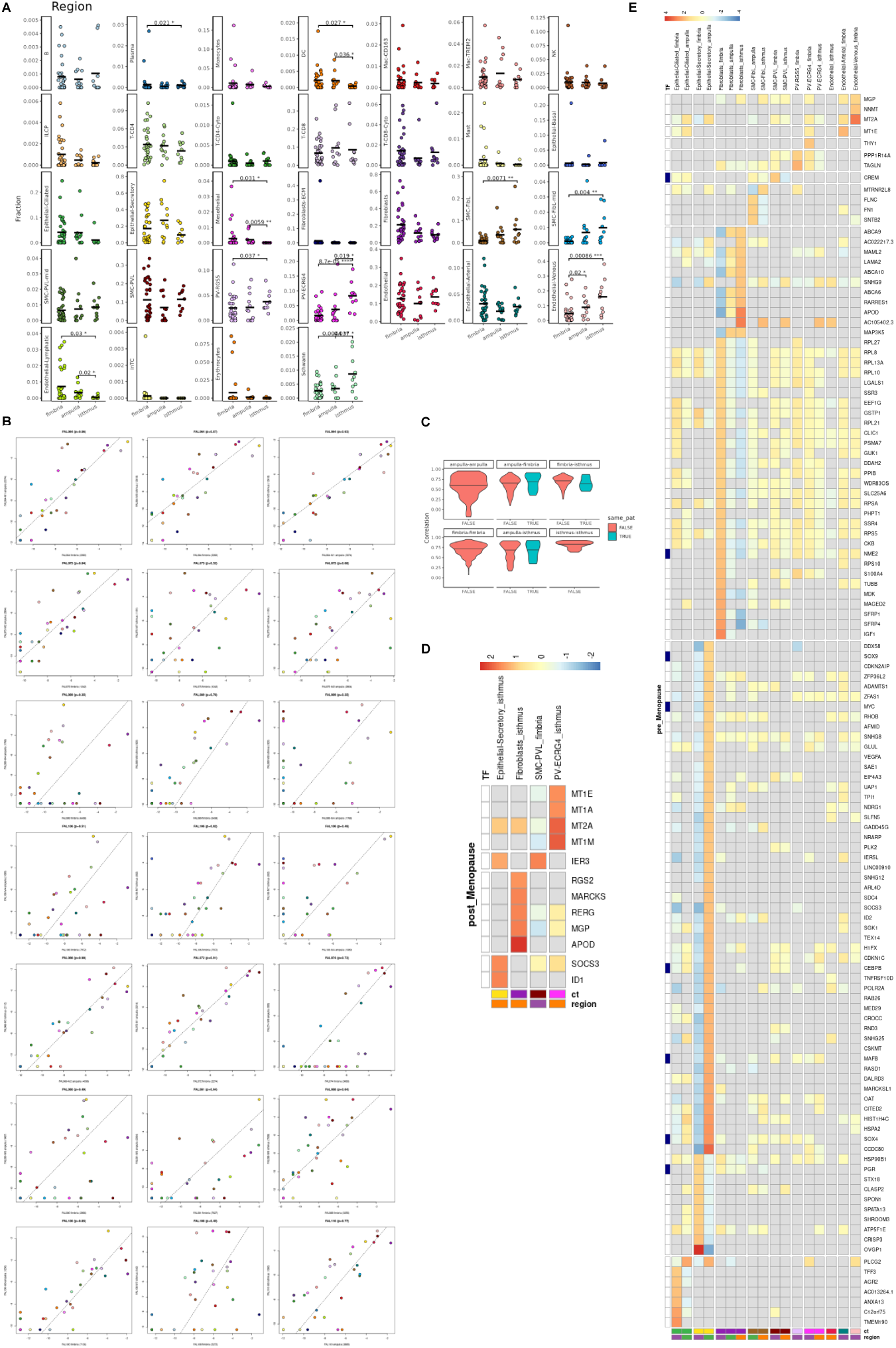
Regional specification in the fallopian tube. **a,** Cell-type abundance comparison across sample by FT region. *P* values by pairwise Mann-Whitney test within each cell-type are shown. Only PV-ECRG4 and Endothelial-Venous cells showed statistically significant difference by FDR corrected Kruskal-Wallis test across all cell-types (see also Fig. 2H). **b,** Comparing samples from different regions of the same subject. Intra-subject cell-type abundance comparison across different regions. Units are cell-type abundance (log2 fraction), axis labels show the region and the number of cells, Pearson correlation shown above each plot. **c,** Distributions of Pearson correlations between cell-type abundances within or across regions, within the same subject or across different subjects. **d**, Controlled DEGs in pre-menopausal samples across the different FT regions, showing LFC by genes (rows, marking transcription factors on the left), and cell-types and FT region (columns). **e,** Controlled DEGs in post-menopausal samples.

**Supplementary Figure 11:**
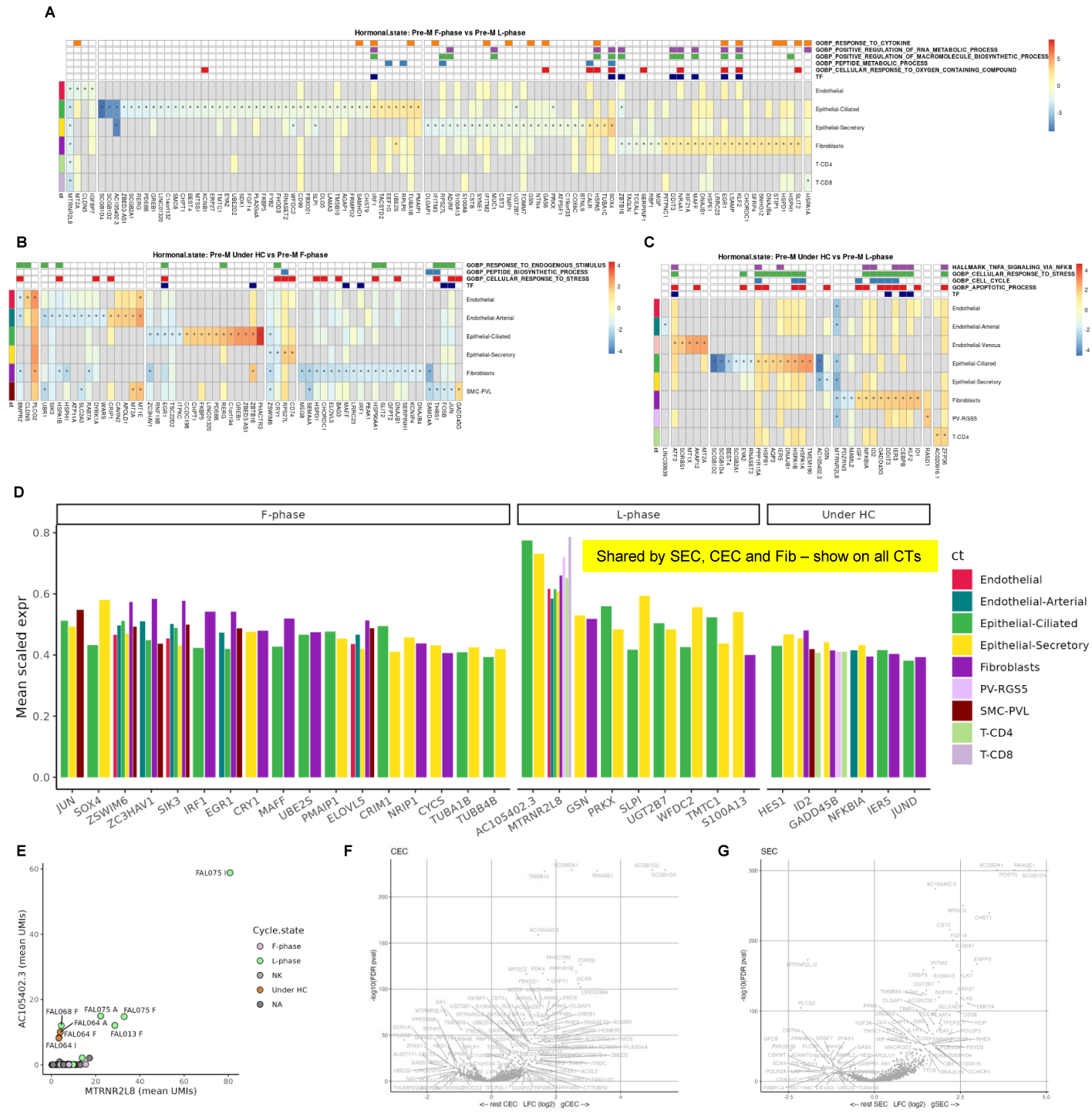
Menstrual cycle and hormonal contraception (HC) influence in the fallopian tube. **a,** Controlled DEGs of pre-menopausal follicular vs luteal-phase samples, showing LFC by genes (columns), marking transcription factors and assignment to GOBP (Gene-Ontology Biology Process) enriched gene sets on top, and cell-types (rows). Asterisks mark absolute LFC ≥ 1. **b,** Comparison of pre-menopausal samples using HC vs Follicular-phase samples. **c,** Comparison of pre-menopausal samples using HC vs Luteal-phase samples. **d,** Genes enriched in each menstrual cycle state. Showing genes shared by SECs, CECs and fibroblast. Showing mean scaled expression (as in Fig. 2C ternary plots) per cell-type (including other cell-types where the gene was differentially expressed), filtering genes with at least 0.4 scaled expression. **e,** Mean UMIs per sample of *AC105402.3* and *MTRNR2L8* genes, coloured by menstrual cycle state, subject and FT region shown for top samples. **f,** DEGs comparing gCECs with remainder of CECs. **g,** DEGs comparing gSECs with remainder of CECs.

**Supplementary Figure 12:**
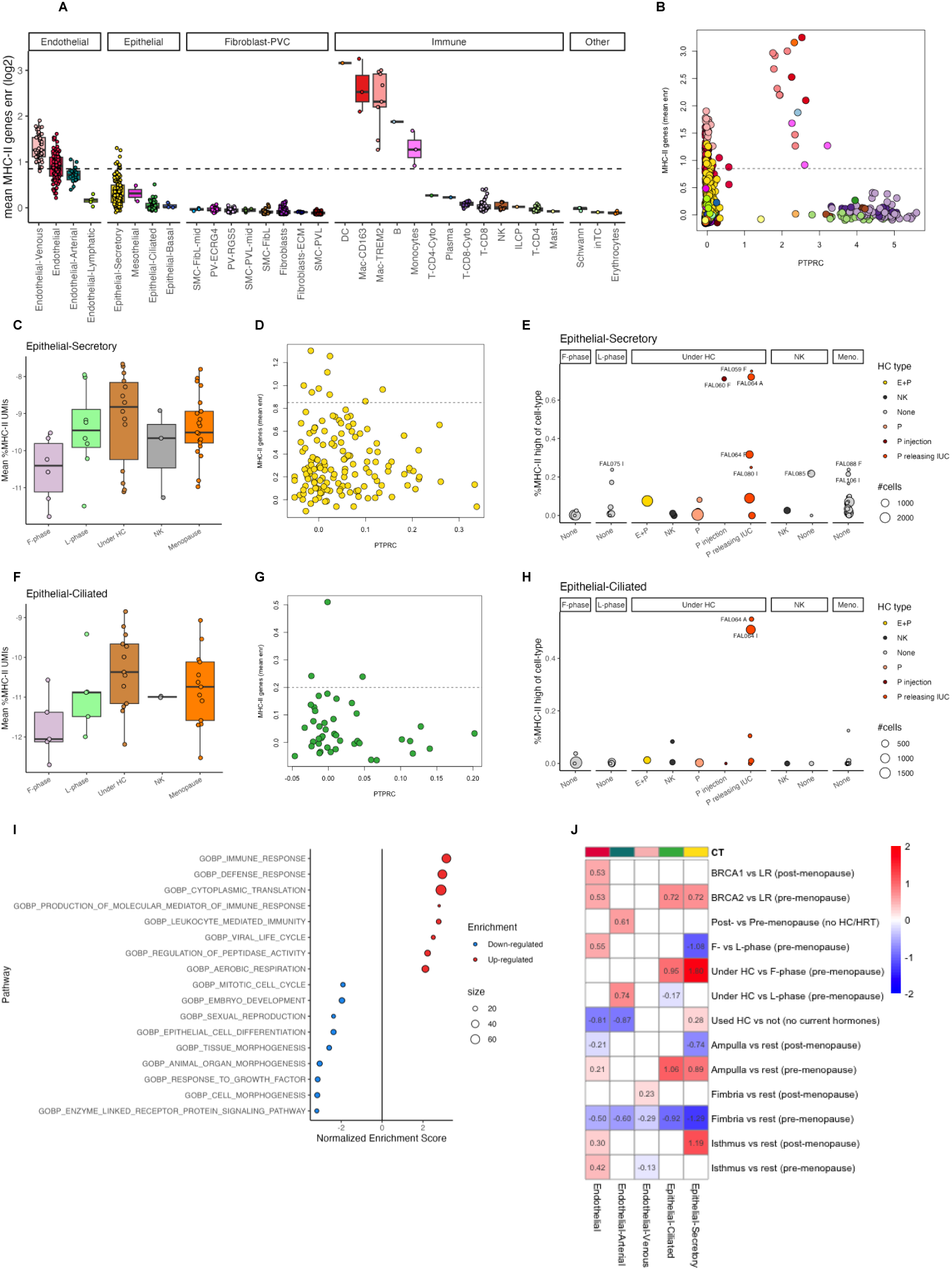
MHC-II expression across cell types and hormonal regulation. **a,** Mean MHC-II genes enrichment per metacell by cell-type and broader cell group, dashed line marks minimum value in APC metacells. **b,** Mean MHC-II genes enrichment by *PTPRC* (which encodes the immune marker CD45), showing the MHC-II high venous endothelial cells and SECs are not endothelial / SECs – immune doublets. **c,** Fraction of MHC-II genes UMIs per sample by menstrual cycle state in SECs. **d,** Fraction of MHC-II genes in SECs, dashed line marks cutoff defining MHC-II high SECs. **e,** Fraction of MHC-II high SECs per sample by menstrual cycle state and type of HC. **f,** Fraction of MHC-II genes UMIs per sample by menstrual cycle state in CECs **g,** Fraction of MHC-II genes in CECs, dashed line marks cutoff defining MHC-II high CECs. **h,** Fraction of MHC-II high CECs per sample by menstrual cycle state and type of HC. **i,** Gene set enrichment analysis of MHC-II high SECs vs rest of SECs, using Gene Ontology Biological Process (GOBP) **j,** Mean controlled differential expression of MHC-II genes across all checked conditions. showing subjects where all MHC-II genes were co-ordinately enriched or depleted.

**Supplementary Figure 13:**
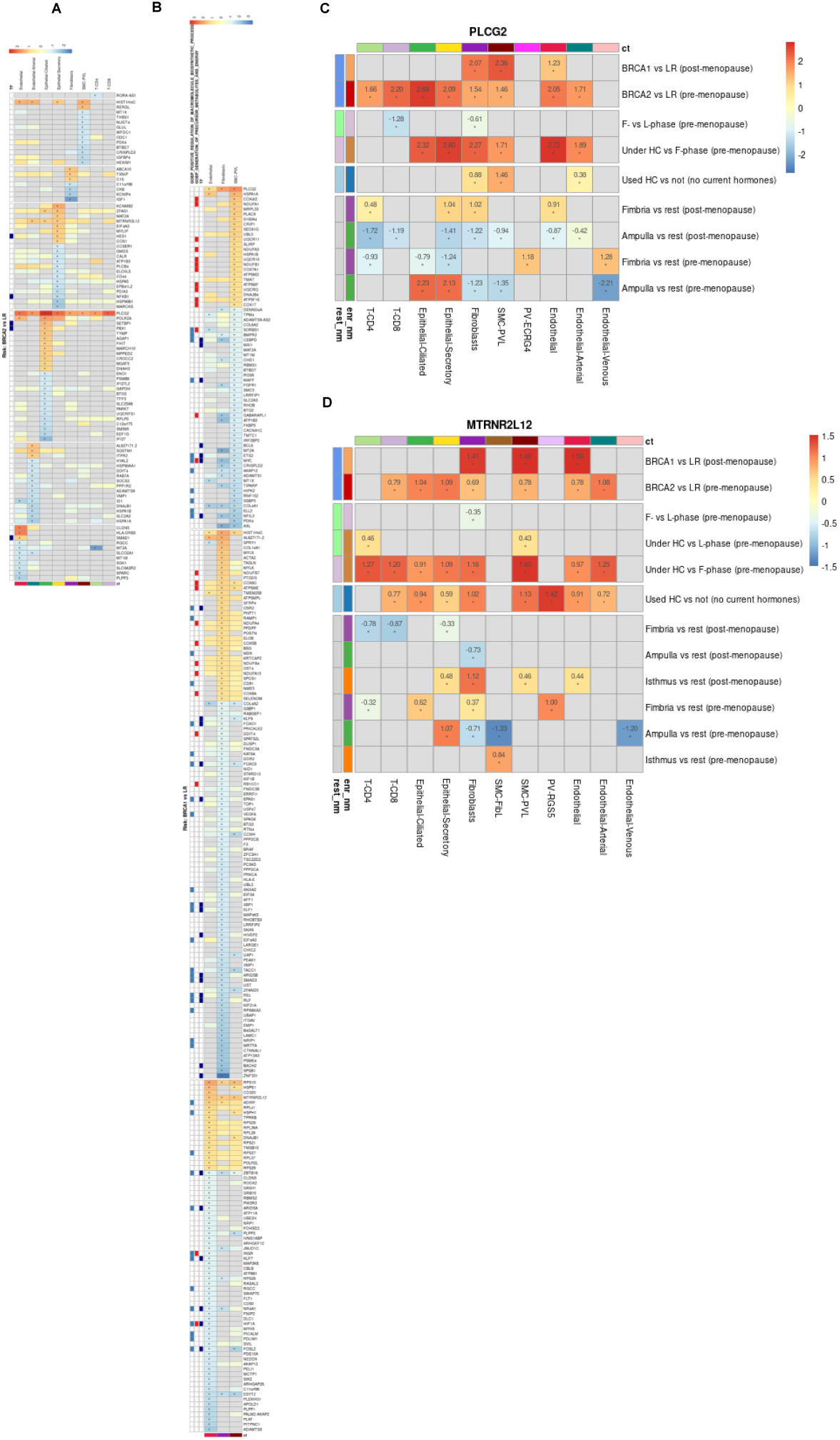
BRCA1/2-associated genes per metacell. **a,** Heatmap comparing samples from pre-menopausal *BRCA2* vs low-risk (LR) subjects, showing genes with absolute LFC ≥ 1 per metacell cell type. TF: transcription factor **b,** Heatmap comparing samples from pre-menopausal *BRCA1* vs low-risk (LR) subjects, showing genes with absolute LFC ≥ 1 per metacell cell type. TF: transcription factor **c,** All cell-type and differential expression (DE) analysis runs where *PLCG2* had an absolute LFC ≥ 1. **d,** All cell-type and DE analysis runs where *MTRNR2L12* had an absolute LFC ≥ 1.

**Supplementary Figure 14:**
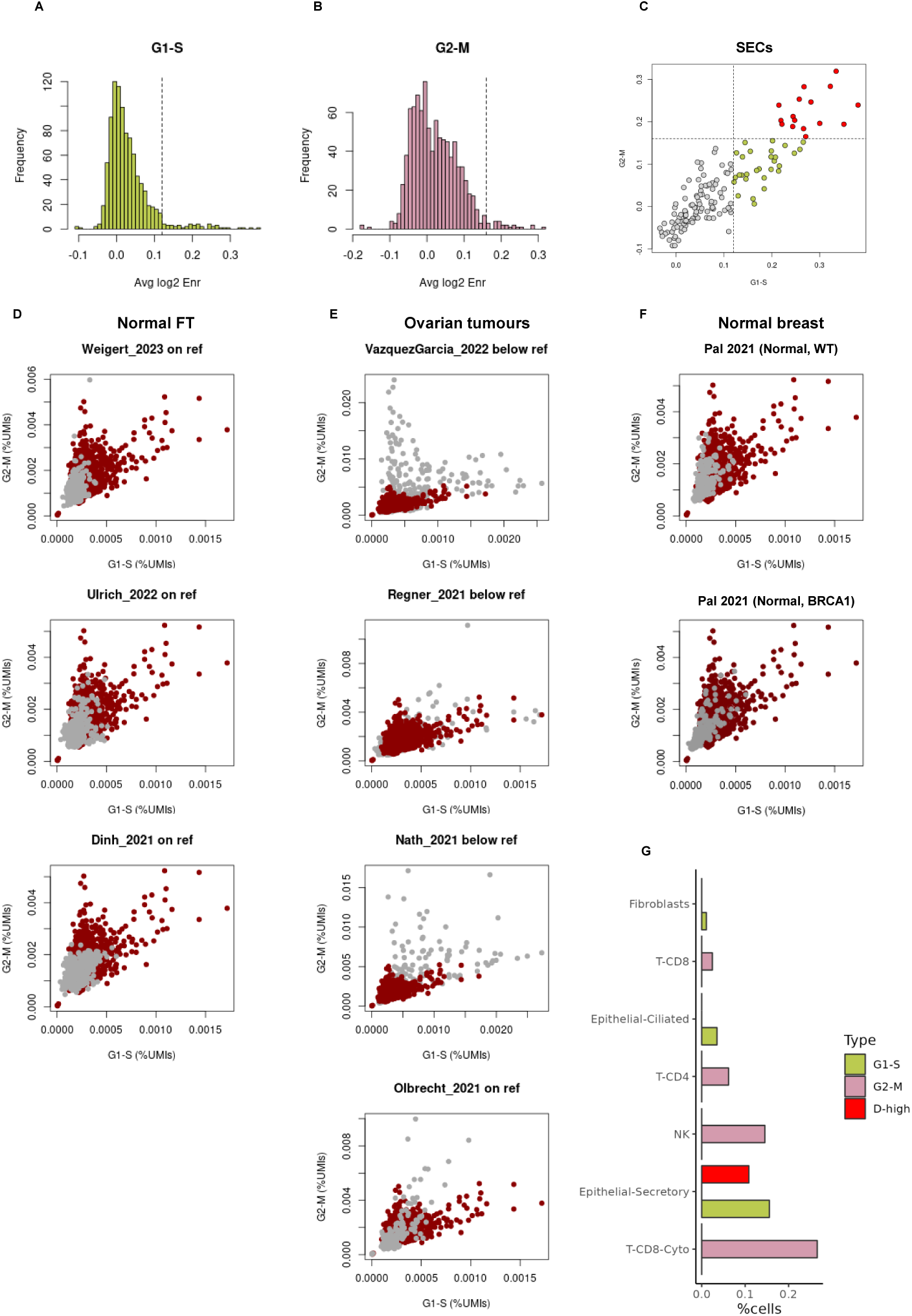
G1–S and G2–M core gene distribution across published datasets. **a,** Distribution of G1–S core genes (from Dominguez et. al 2016 ^33^) mean enrichment per metacell, dashed line marks the cutoff for G1–S ^+^ metacells **b,** Distribution of G2–M core genes **c,** G1–S vs G2–M core genes mean enrichment per SEC metacell, colour by G1–S / G2–M cutoffs (colour legend at panel g) **d,** Fraction of G1–S and G2–M core gene UMIs per metacell in our dataset (red) vs other normal FT scRNA-seq datasets (grey) **e,** As shown in d, but for ovarian cancer datasets **f,** As shown in d, but for a normal breast dataset **g,** Frequency of G1–S ^+^, G2–M ^+^ and G1–S ^+^-G2–M ^+^ (D-high) cells per cell-type, only showing cell-type with cells in any of these categories.

**Supplementary Figure 15:**
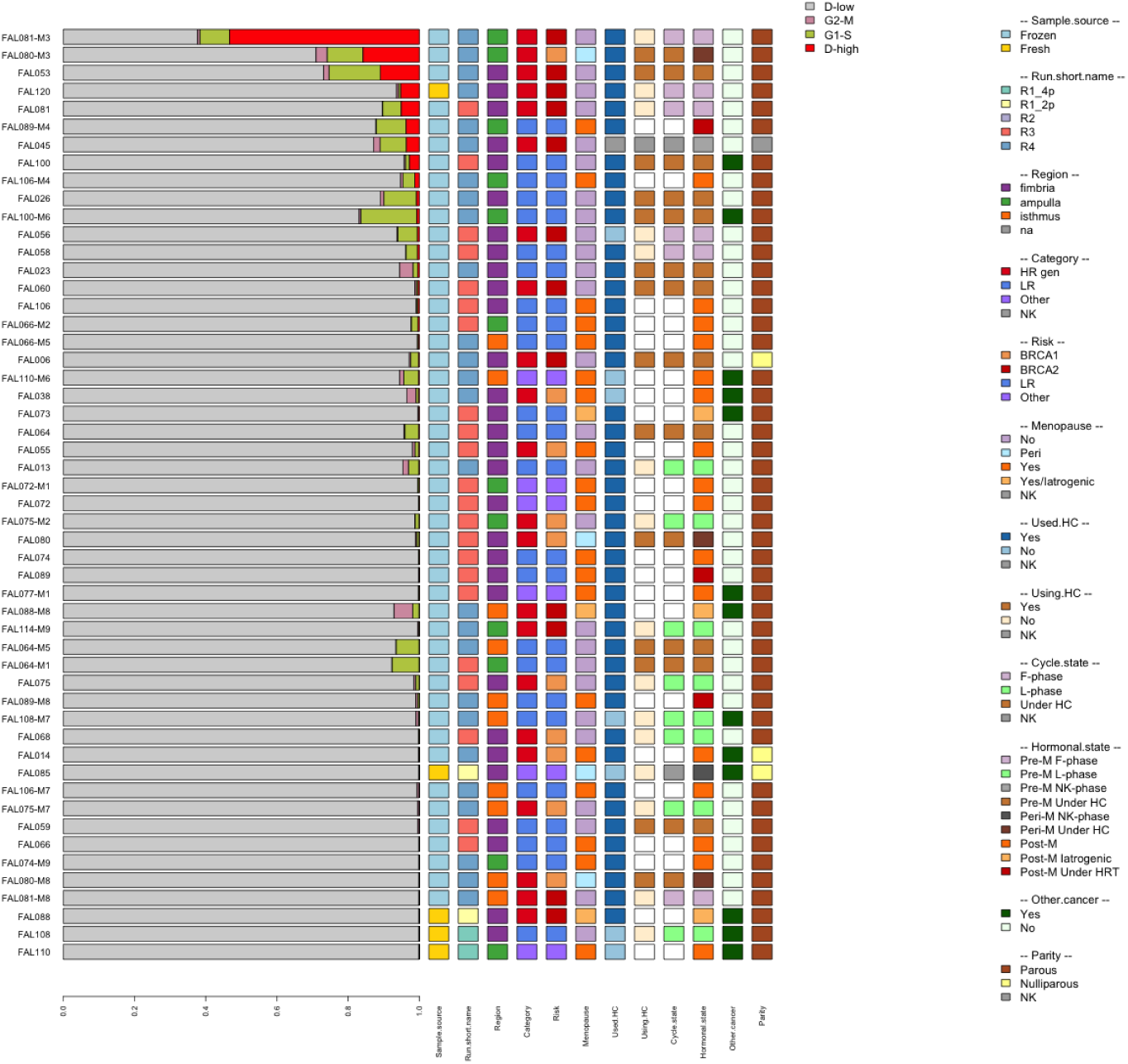
Sample composition based on expression of G1–S and G2–M core genes. **a,** Sample composition by expression of G1–S and G2–M core genes, showing samples with at least 100 cells, samples ordered by decreasing frequency of cells with double-positivity for G1–S and G2–M genes; samples clinical features shown to the right.

**Supplementary Figure 16:**
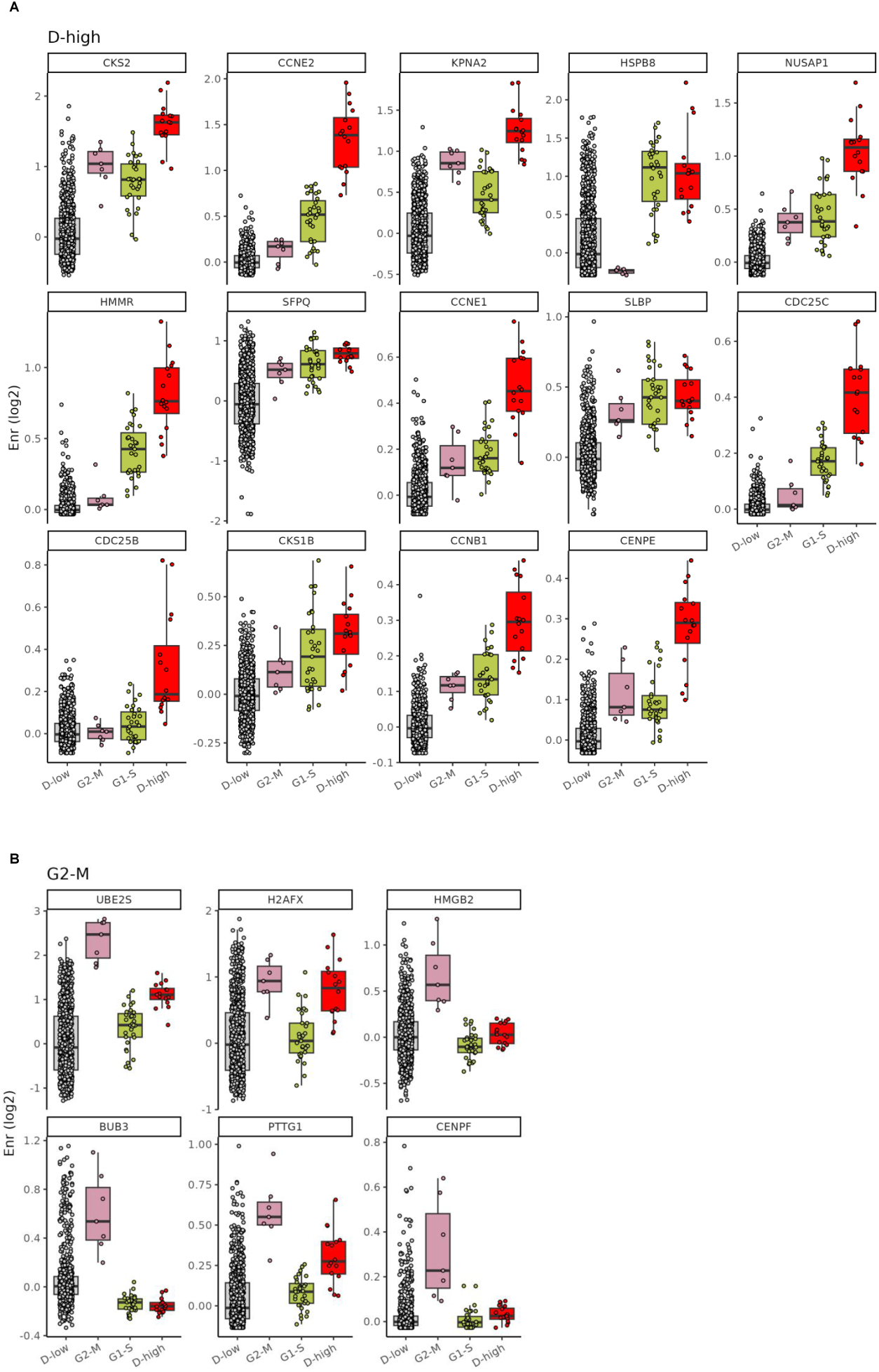
Gene enrichment across G1–S/G2–M categories. **a,** Core G1–S and G2–M genes enrichment across the G1–S / G2–M categories for genes with highest mean enrichment in the G1–S ^+^-G2–M ^+^ (D-high) group, genes ordered by mean enrichment, only showing genes with mean enrichment above 0.25. **b,** As shown in a, but for genes with highest mean enrichment in the G2–M**+** group (there are no core genes with highest mean enrichment in the G1–S ^+^ group).

**Supplementary Figure 17:**
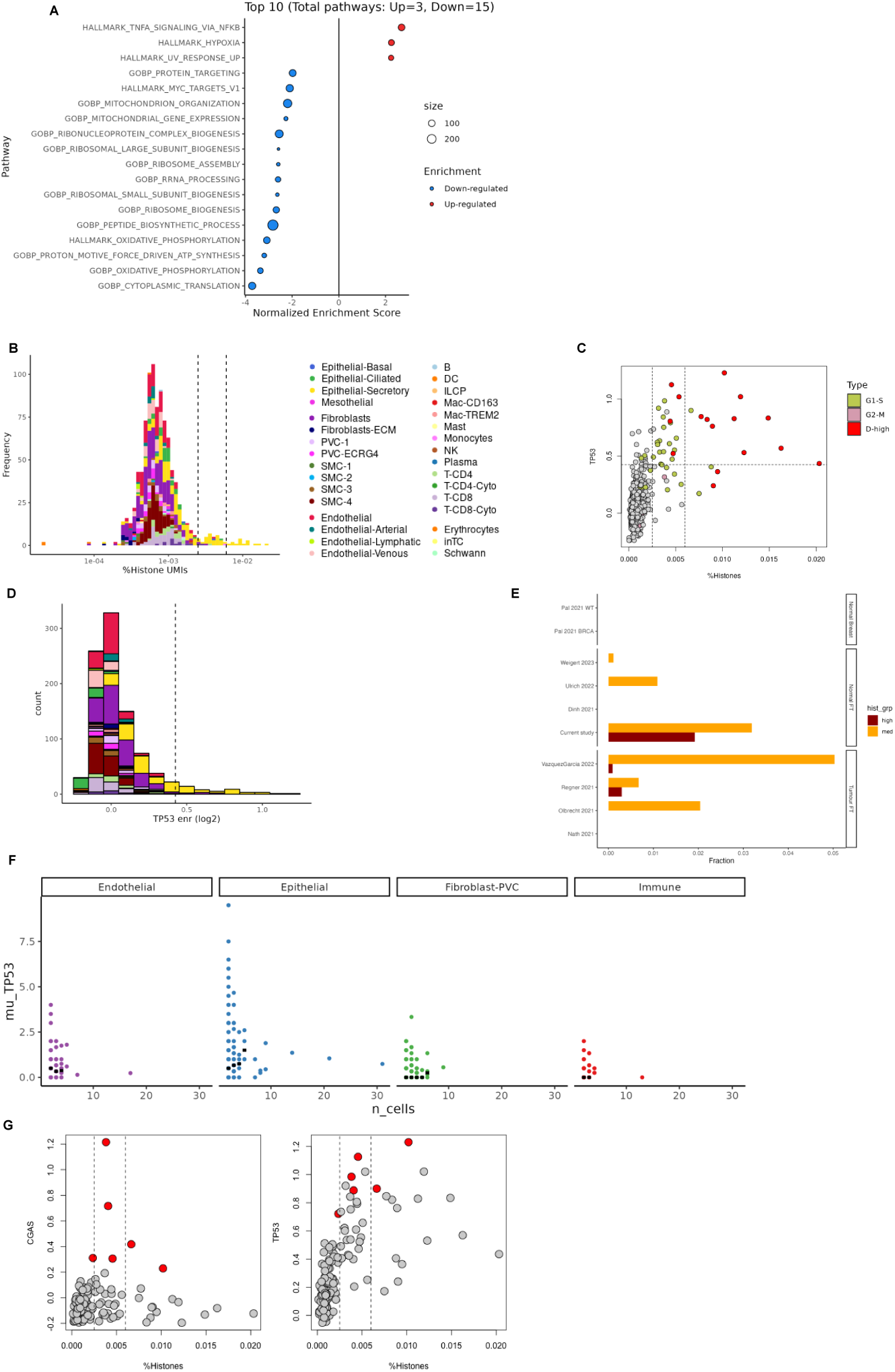
SEC metacell distribution across expression levels of TP53 and histone genes. **a,** Gene set enrichment analysis of DEGs between the G1–S ^+^-G2–M ^+^ (D-high) vs the G1–S ^−^-G2–M ^−^ (D-low) cells. **b,** Histogram of fraction of UMIs from histone coding genes (prefixed HIST#H) per metacell, bars coloured by their metacell cell-type composition, and dashed lines delineate metacells to low, mid and high-histone groups. **c,** *TP53* gene enrichment vs mean histone UMIs per metacell, dashed lines mark *TP53* and histone grouping cutoffs, metacells coloured by G1–S/G2–M groups. **d,** Histogram of metacells *TP53* gene enrichment, bars coloured by their metacell cell-type, dashed line delineates metacells to low- and high-*TP53* groups. **e,** Fraction of histone-med and histone-high cells in related datasets, stratifying cells by the same histone level cutoffs as in c, showing normal FT and breast datasets and HGSOC datasets. **f,** *TP53* mean expression in clone cells by clone size per SNV clone, grouping by high-level cell-type, black ticks mark median value for clone sizes. **g,** *CGAS* and *TP53* gene enrichment vs mean histone UMIs per SEC metacell; dashed lines mark histone grouping cutoffs; high CGAS metacells marked in red.

**Supplementary Figure 18:**
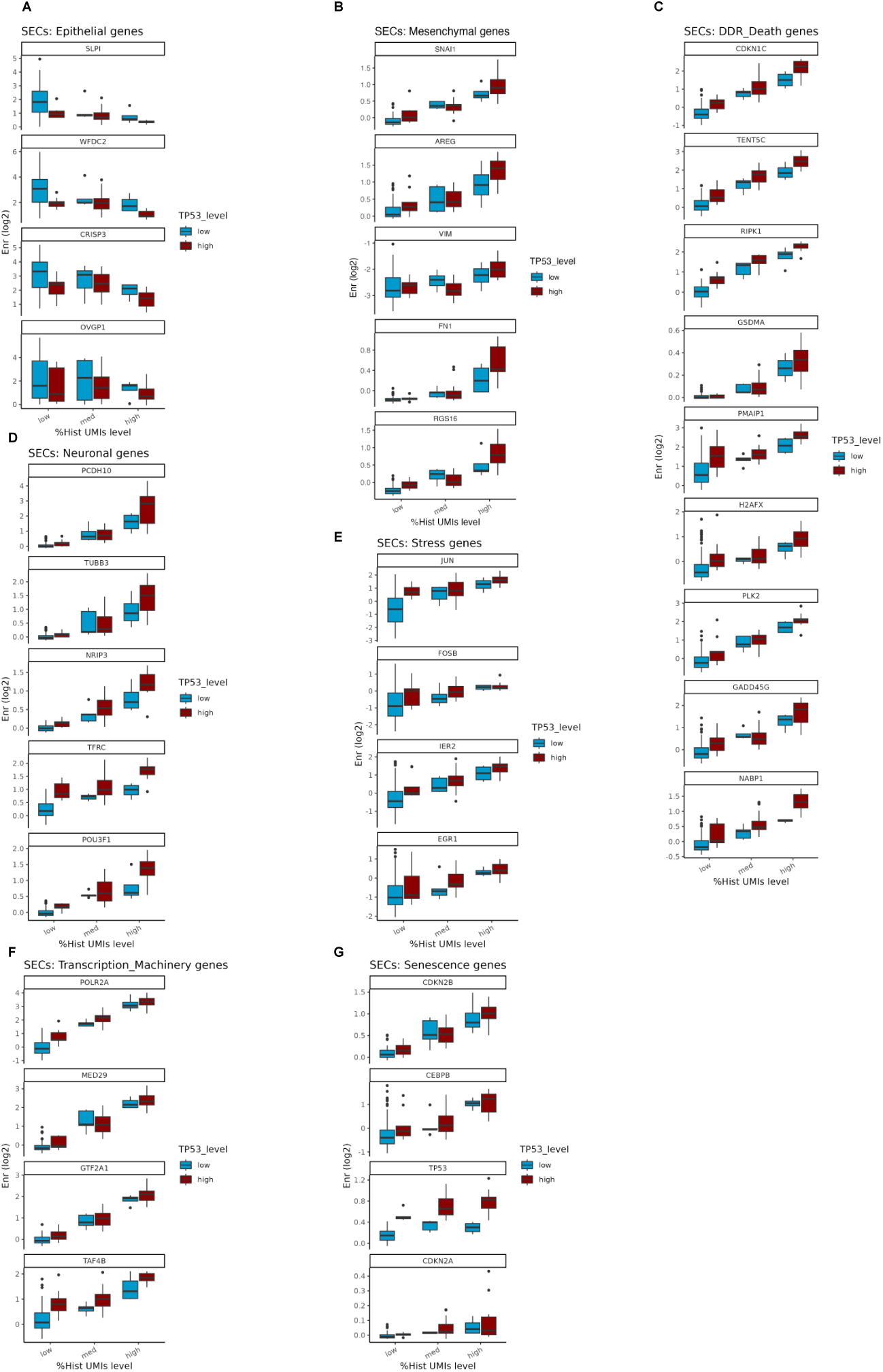
SEC metacells gene enrichment of several key genes across the Histone / *TP53* groups. **a,** Epithelial cells. **b,** Mesenchymal cells. **c,** DNA damage response and cell death genes. **d,** Neuronal genes. **e,** Stress genes. **f,** Transcriptional machinery genes. **g,** Senescence genes.

**Supplementary Figure 19:**
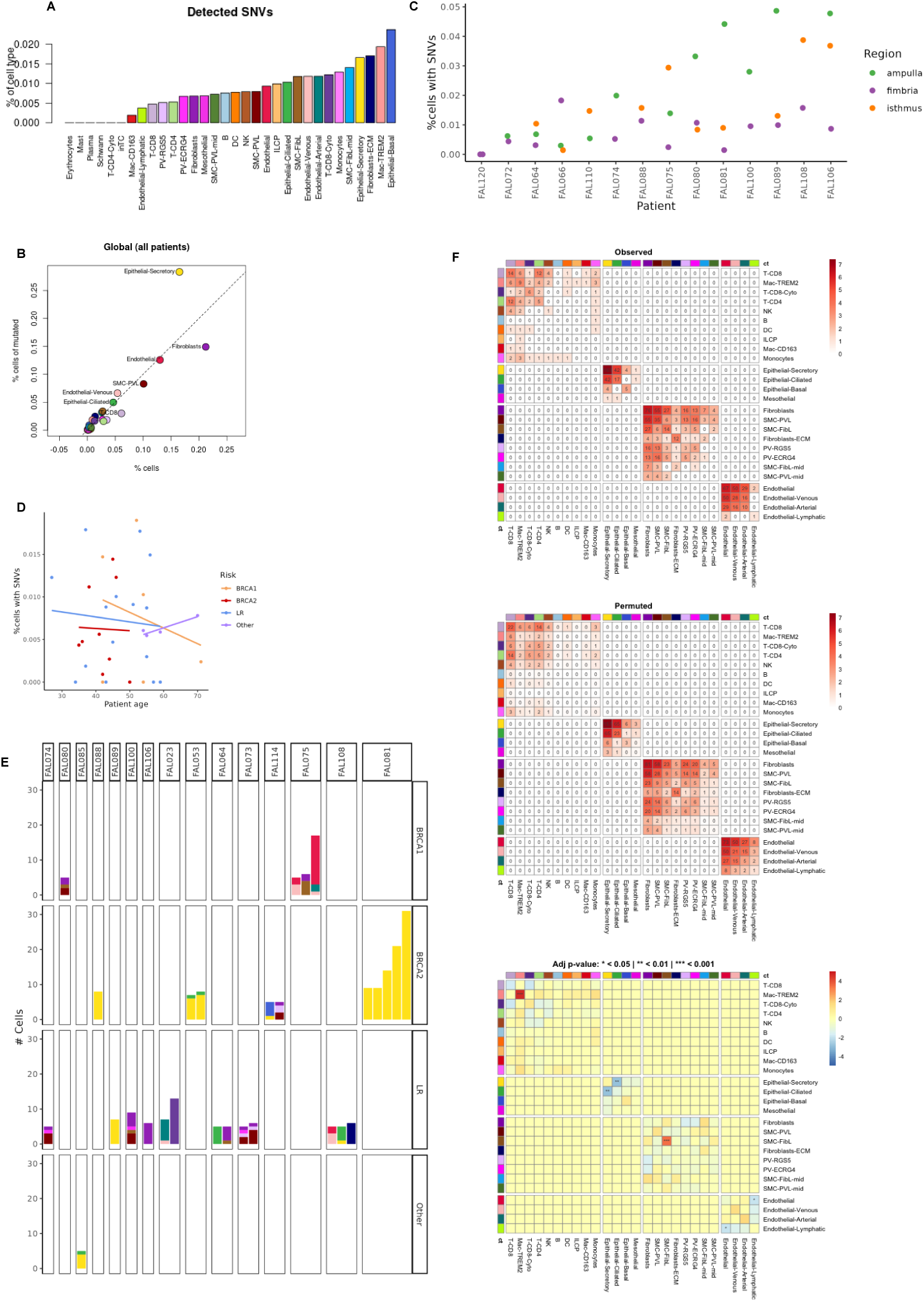
SNV distribution across fallopian tube regions, metacells and risk categories. **a,** SNV frequency per cell-type. **b,** Fraction of cell-type of all cells and of cells with a detected SNV, showing the higher SNV contribution of SECs. **c,** SNV frequency per subject and region (for multi-region subjects), subjects ordered by mean SNV frequency **d,** Fraction of cells with a detected SNV by sample versus subject age, coloured by risk, with linear fitted lines per risk group. **e,** Cell-type composition of large (≥ 5 cells) SNV clones by subject and risk category. **f,** SNV sharing across cell-types: showing the observed number of distinct shared SNVs per pair of cell-types on top, the expected number by random after permuting cell-type labels (hence maintaining the same cell-type frequencies and clone sizes) and the z-score enrichment of the observed over the expected by random (bottom).

**Supplementary Figure 20:**
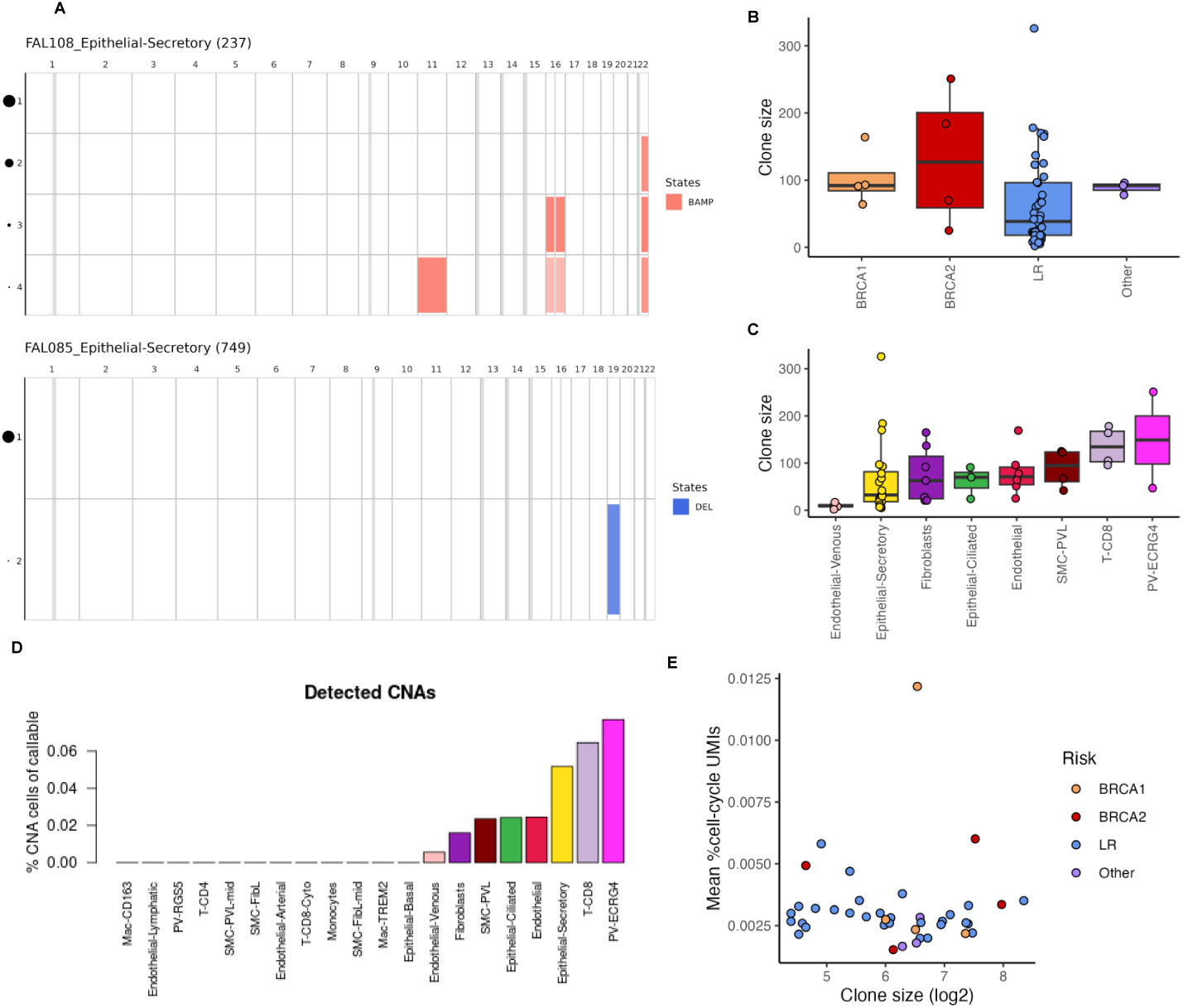
Distribution of inferred SCNA clones across risk categories and meta cells. **a,** Representative plots of SCNA clones in SECs for two subjects, number of cells in parenthesis. SCNAs are marked by colour, with chromosomes in columns and clones in rows. Clone 1 is the normal diploid one, circle size by clone number represents the clone size. **b,** SCNA clone size by risk category. **c,** SCNA clone size by risk category by cell-type. **d,** SCNA frequency by cell-type. **e,** Mean fraction of UMIs from cell-cycle gene per SCNA clone by clone size, colouring clones by risk.

**Supplementary Figure 21:**
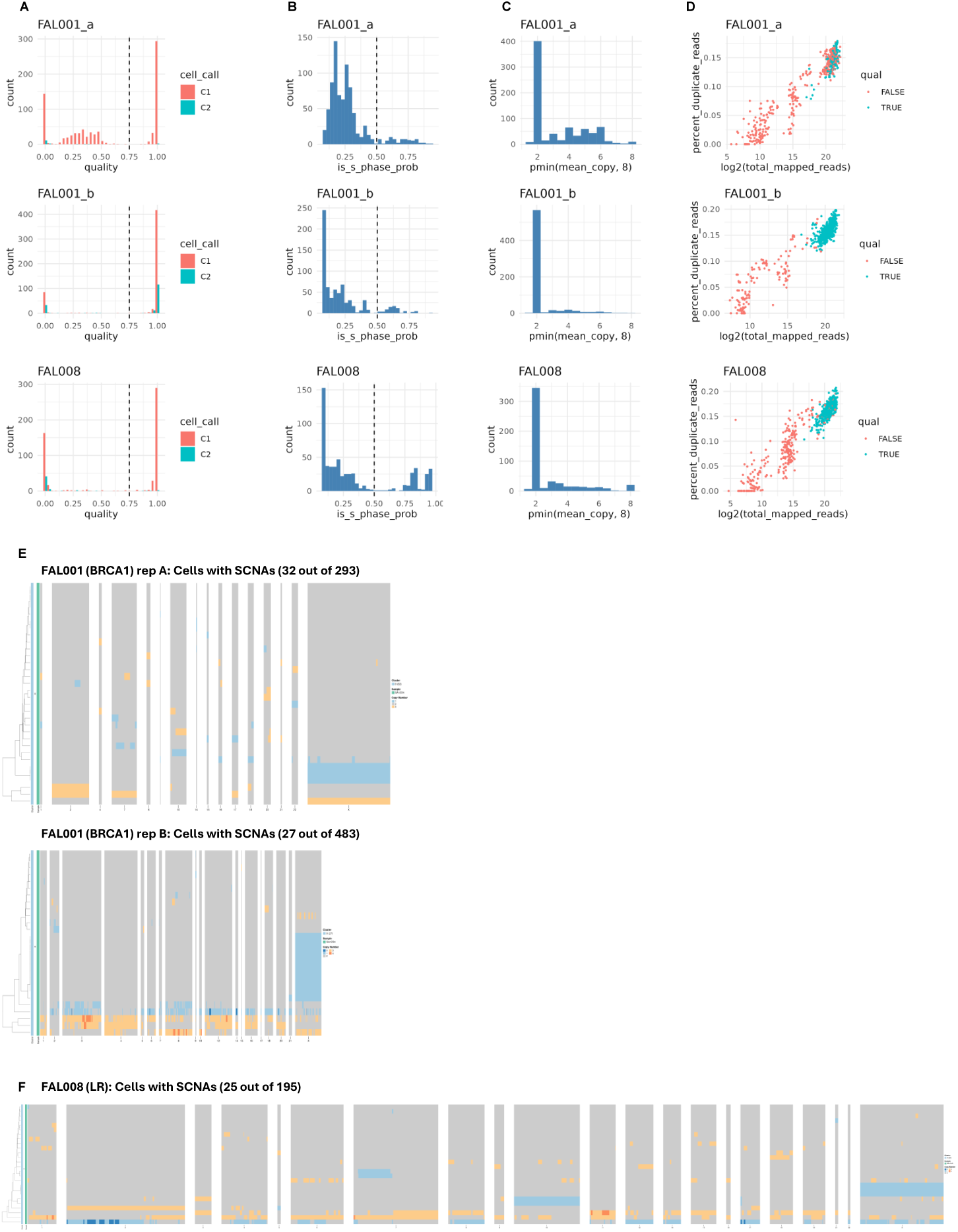
Copy-number scDNA-seq profiles of a low-risk and *BRCA1* subject. **a,** Distribution of cells quality by cell call (C1 mark valid cells). **b,** Probability of being in S-phase, dashed line mark cutoff for filtering suspected S-phase cells. **c,** Mean copy-number per cell, showing that most of the cells are normal diploids. **d,** Percentage of duplicate reads vs total number of mapped reads, cells that passed quality cutoff shown in cyan. **e,** Copy number profile of only the non-diploid cells (rows) and the non-diploid genomic bins (columns) for FAL001 (2 biological replicates). **f,** Copy number profile of only the non-diploid cells (rows) and the non-diploid genomic bins (columns) for FAL008 sample.

**Supplementary Figure 22:**
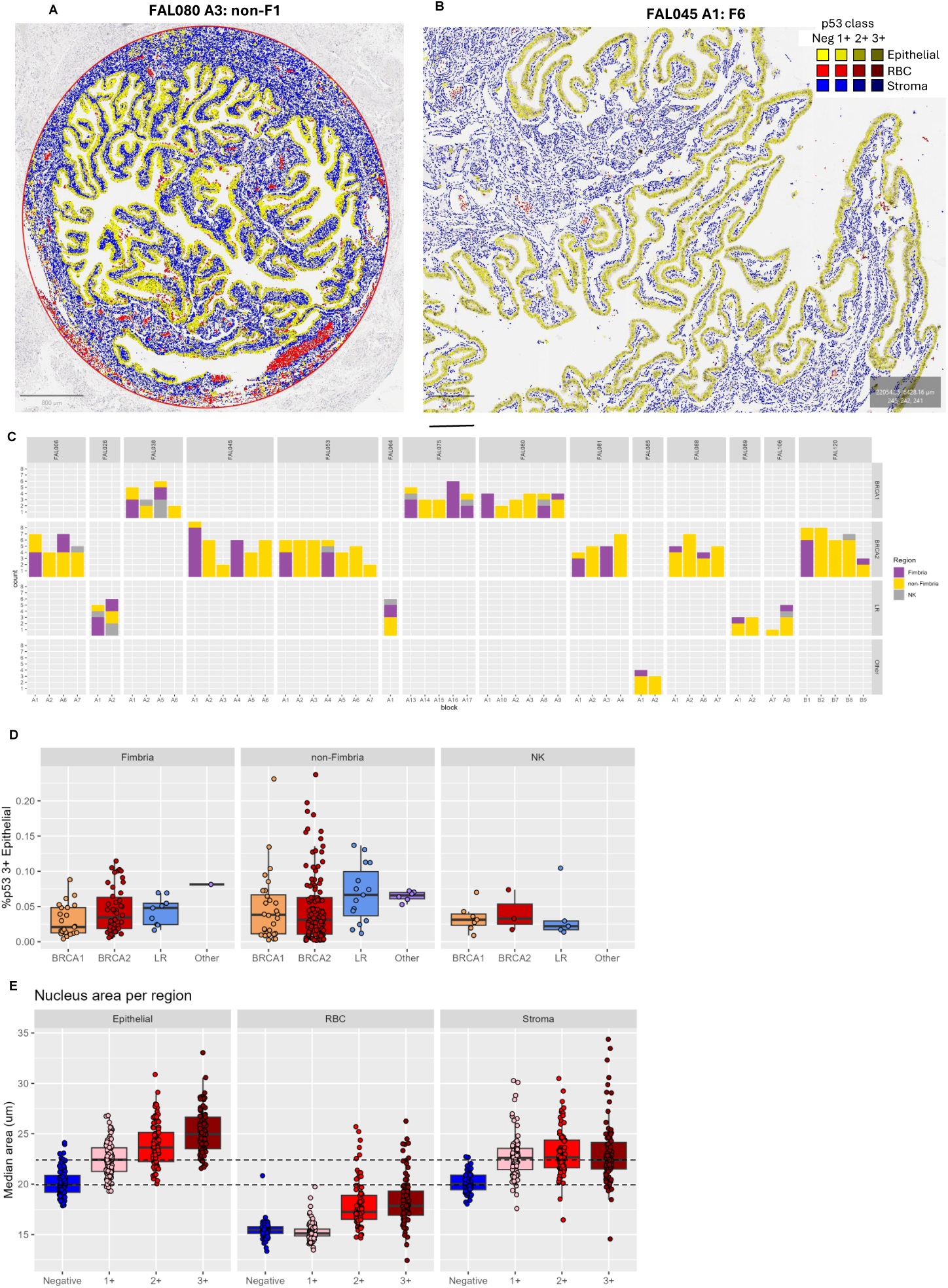
Automated p53 IHC analysis in fallopian tube tissues. **a,** Example of a p53 IHC slide overview; scale bar length: 800 µm, showing a whole non-fimbrial region in FAL080 slide A3, cells coloured by their classification to cell-type (Epithelial, Stroma, RBC) and p53 positivity (Neg, 1+, 2+, 3+), as shown in the legend on the right. **b,** Example of a p53 IHC slide; scale bar length: 250 µm, from a large fimbrial region in FAL045 slide A1. **c,** Number and classification (fimbrial, non-fimbrial, NK) of regions per slide and subject, separating subjects by risk (rows). **d,** Fraction of Epithelial 3+ (strong p53 positive) cells of all epithelial cells per region, grouping by risk and stratifying by FT region. **e,** Median nuclei area per region, grouping by p53 state and stratifying by cell-type, dashed lines at median values of Epithelial Neg and 1+ groups.

**Supplementary Figure 23:**
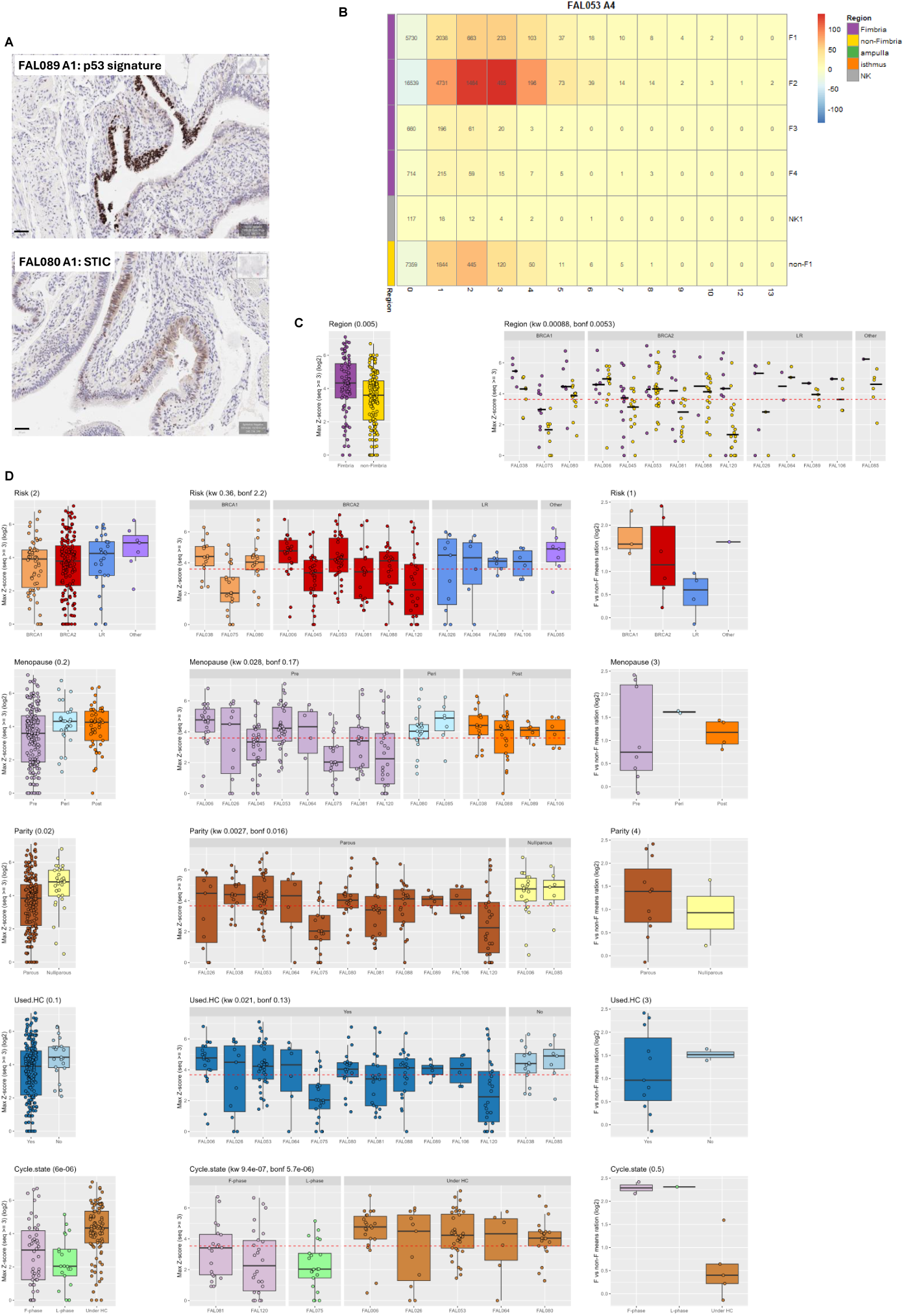
Automated detection of p53 signature/STIC in fallopian tube tissues. **a,** Raw p53 IHC staining focused on a p53 signature in FAL089 slide A1 (top) and a STIC in FAL080 slide A1 (bottom); scale bar length: 50 µm. **b,** Heatmap showing Epithelial 3+ streak length enrichment per region and streak length for FAL053 slide A4. Number shows the observed number of cells with a particular streak length (columns, capped at 13) in a region (rows). Cell colours are the Z-score of that observed number over the expected one by random considering the frequency of Epithelial 3+ cells in the region. **c,** Fimbrial streak enrichment score. Showing the log2 ratio between the mean streak length score in the fimbria vs non-fimbria regions per subject, grouping by the different clinical features, *P* values by Bonferroni corrected Kruskal-Wallis tests over all clinical features shown on top. Streak enrichment score (maximal Z-score of streaks length 3 or more) per region in slide. Grouping from top by FT region, risk, menopausal status, parity, HC usage history and menstrual cycle state. P-values by Bonferroni corrected Kruskal-Wallis tests over all clinical features shown on top. **d,** Streak enrichment score (maximal Z-score of streaks length 3 or more) per region in slide. Grouping from top by FT region, risk, menopausal status, parity, HC usage history and menstrual cycle state. P-values by Bonferroni corrected Kruskal-Wallis tests over all clinical features shown on top.

**Supplementary Figure 24:**
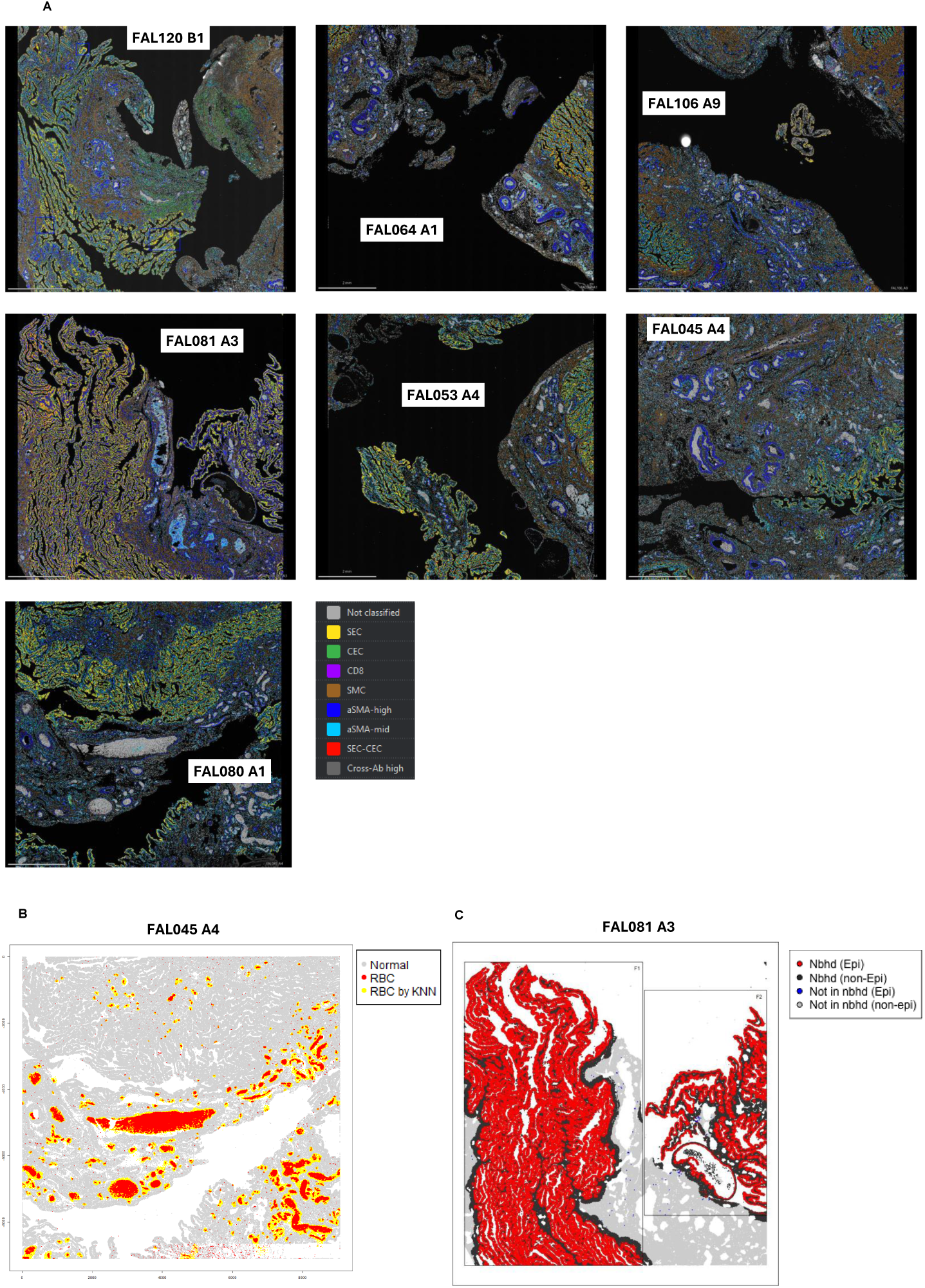
Protein-inferred spatial cellular distribution in the fallopian tube. **a,** Whole slide view of the 7 COMET slides analysed; scale bar length: 2mm, cells coloured by classification, with colour legend on the bottom right. **b,** RBC detection in FAL045 slide A4. Red dots mark the initial set of cells identified as RBCs by the filtering criteria, yellow dots mark cells further classified as RBC by iterative (200 iterations) majority vote on their k-NNs (K = 25). **c,** Example of an epithelial neighbourhood demarcation. Within each manually marked region (F1 and F2 rectangles in this example) cells are included in the neighbourhood if at least 100 of their 750 k-NNs are epithelial cells. Showing epithelial and non-epithelial cells inside and outside the neighbourhoods.

**Supplementary Figure 25:**
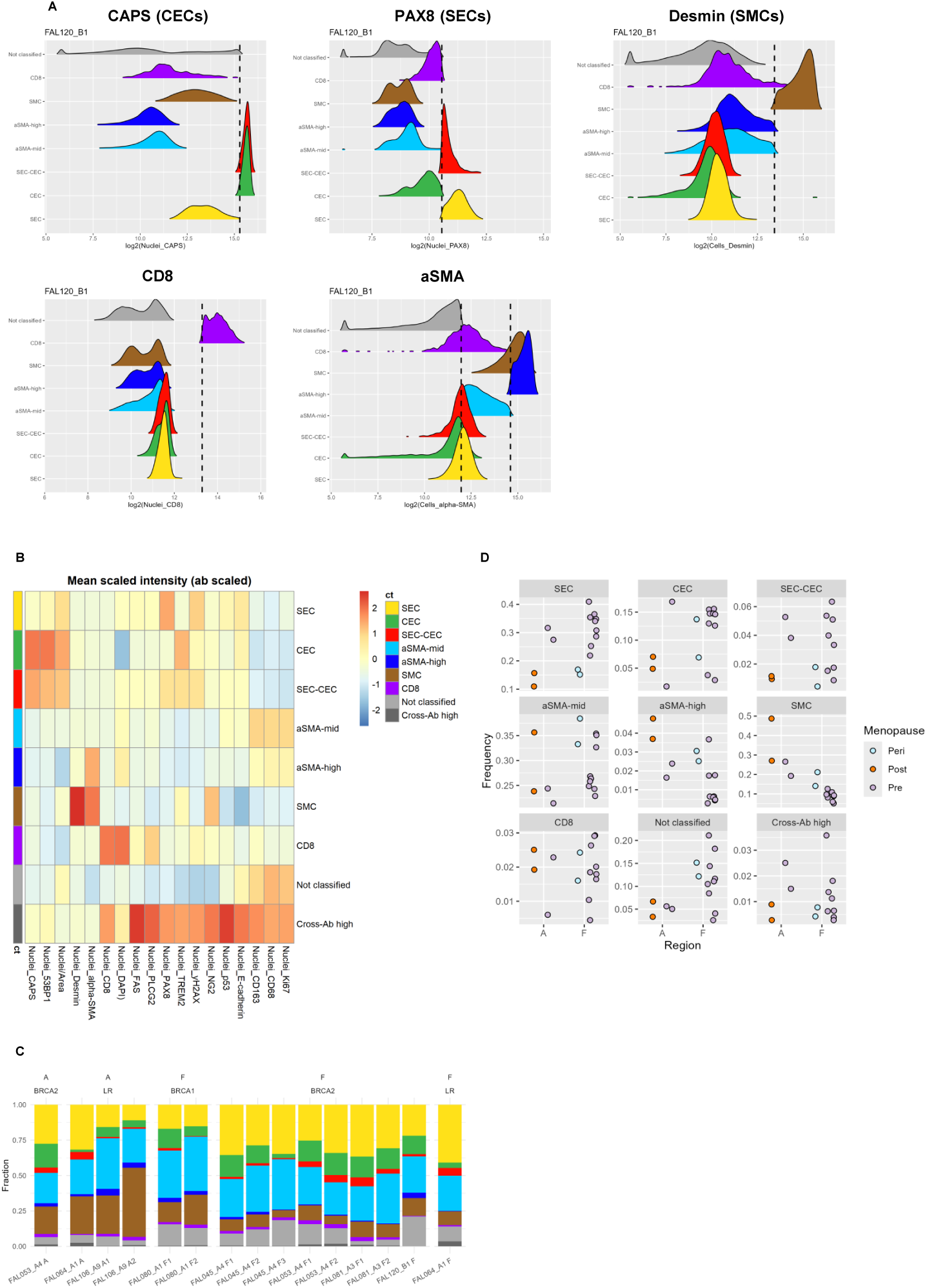
Protein-inferred fallopian tube cellular distribution across region, menopausal status and risk categories. **a,** Example of COMET antibodies used for cell-type classification in slide B1 of FAL120. Each panel shows log2 intensity distribution of an antibody across the different cell-types. Title shows the antibody and its associated cell-type, with dashed line marking the cell-type classification cutoffs for that cell-type. **b,** Mean antibody intensity (scaled per antibody) across all cells by cell-type. **c,** Cell-type composition of the different tissue fragments across all slides, grouped by risk and FT region. **d,** Cell-type abundance in epithelial neighbourhoods grouped by subject menopausal state and stratified by FT region (fimbria and ampulla).

**Supplementary Figure 26:**
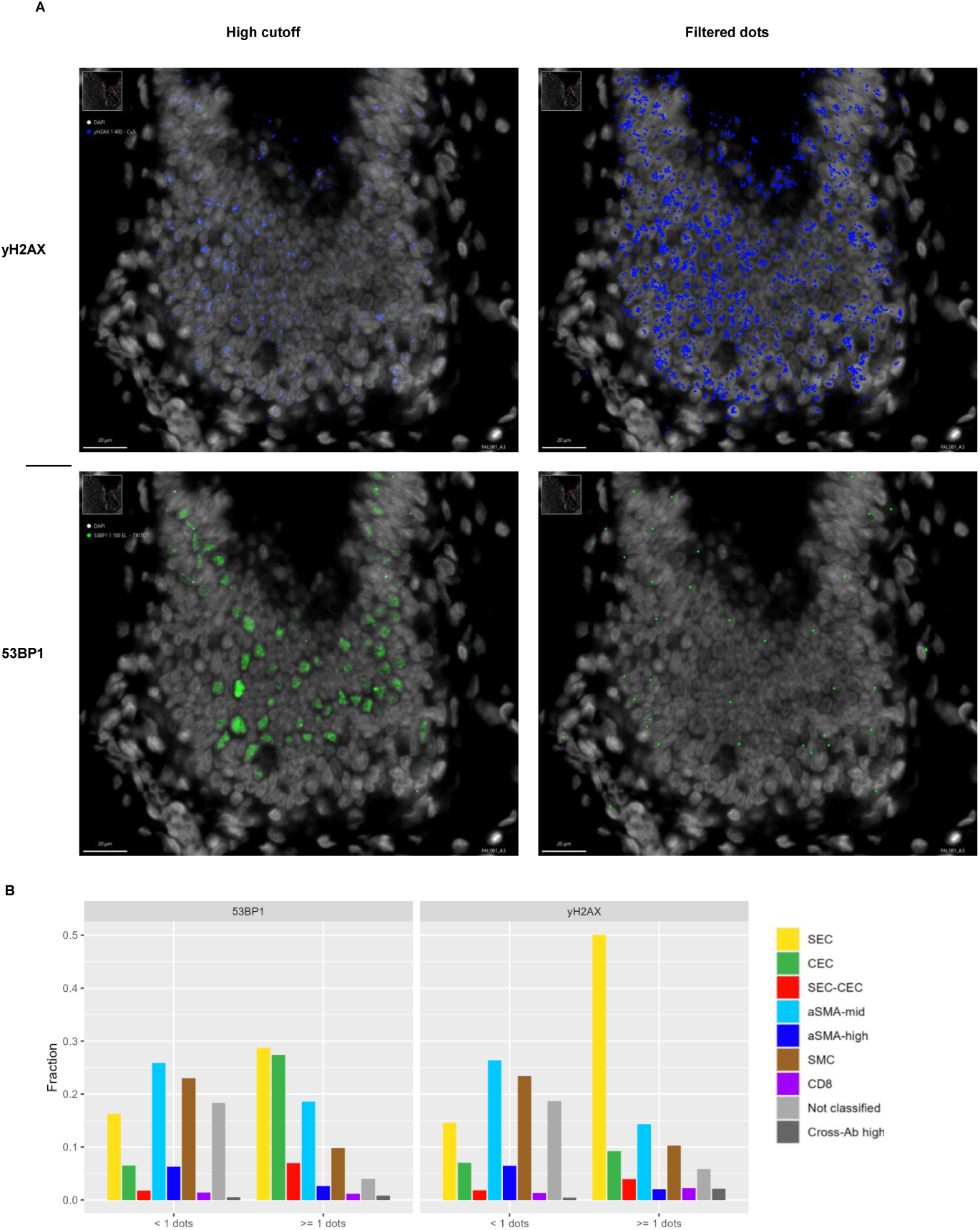
Distribution of γ-H2AX and 53BP1 positivity across cell types. **a,** High-intensity γ-H2AX and 53BP1 staining in a focused region; scalebar length 20 µm, with the dots identified by COMET for each antibody. **b,** Cell-type composition of the dot-positive and dot-negative cell population for γ-H2AX and 53BP1

